# BDF6 deficiency severely compromises intracellular amastigote development and infectivity of *Trypanosoma cruzi*

**DOI:** 10.1101/2025.07.21.665854

**Authors:** Victoria Boselli, Virginia G. Perdomo, Florencia Pacini, Juan A. Cueto, Alejandro Pezza, Patricia S. Romano, Roberto Docampo, Ana Rosa Perez, Esteban C. Serra

## Abstract

*Trypanosoma cruzi*, the causative agent of Chagas disease, relies on complex gene regulatory mechanisms to adapt to its diverse host environments. In recent years, it has been established that epigenetics plays an essential role in these mechanisms via the regulation of chromatin structure. Bromodomain-containing factors (BDFs), known for recognizing acetylated lysines on histones, have emerged as key factors in chromatin remodeling complexes. Among the eight predicted BDFs in *T. cruzi*, BDF6 is part of a TINTIN-like complex with MRGx and MRGBP, homologous to components of the NuA4/TIP60 chromatin remodeling complex. We generated knockout (KO) parasites for *bdf6* gene using CRISPR/Cas9 gene editing. BDF6-deficient epimastigotes exhibit normal morphology but decreased size and growth and the resulting metacyclic trypomastigotes displayed drastically reduced infectivity. Strikingly, once inside host cells, BDF6-deficient parasites differentiated into amastigotes but failed to replicate. This intracellular arrest was reversed by episomal complementation of BDF6. Consistently, BDF6-KO parasites also exhibited impaired infectivity in mice, a defect that was also rescued in the add back parasite strains. Our findings highlight BDF6 as a critical regulator of intracellular parasite development, operating in stages beyond epimastigotes where epigenetic plasticity is essential for host adaptation. This striking stage-specific phenotype of BDF6 KO underscores its functional importance and highlights the relevance of epigenetic regulators along *T. cruzi’s* life cycle.

## Introduction

*Trypanosoma cruzi* is the etiological agent of Chagas disease, which affects at least 7 million people globally. Among those infected, 20–30% develop severe clinical manifestations, which means a significant challenge to public health **[1]**. While historically confined to Latin America, migration has facilitated the spread of Chagaś diseases to non-endemic regions, where transmission occurs via blood transfusion, organ transplantation, and congenital routes **[2]**. The limited and often side-effect-prone therapeutic options highlights the urgent need for novel therapeutic targets **[3].**

The life cycle of *T. cruzi* involves both vertebrate and invertebrate hosts, transitioning through extracellular and intracellular stages **[4]**. Such a complex life cycle, involving several morphologically and biochemically distinct parasite forms and different host environments, requires sophisticate regulatory mechanisms to direct gene expression and the adaptive strategies that enable parasites to thrive within distinct hosts. **[5]**. Unlike most eukaryotic cells, *T. cruzi* exhibits polycistronic transcription of protein-coding genes within transcriptional units (PTUs) and lacks canonical promoters or regulatory transcription factors **[6]**. Due to this phenomenon, fine regulation of gene expression occurs through post-transcriptional mechanisms **[7].** Despite this, chromatin remodeling through covalent histone modifications and/or histone variants regulates replication, DNA repair and transcription initiation **[8]**. Many, but not all, of the epigenetic mechanisms that regulate chromatin structure, and consequently these processes, are emerging to be conserved across eukaryotes **[9,10,11].]**. In kinetoplastids, although the field of epigenetics is still emerging, mounting evidence indicates that chromatin regulation, via histone variants, post-translational modifications and three-dimensional genome architecture, plays a potentially crucial role in parasite adaptation and virulence.**[12]**

Bromodomains (BDs) are ∼110 amino acid protein modules that specifically bind to acetylated lysines (AcK) and are considered typical “readers” of histone epigenetic marks. They consist of a four-α-helix bundle (αA, αB, αC, αZ) connected by loops that form a hydrophobic pocket essential for Ac-K recognition **[10,13]**. Upon recognizing Ac-K, BDs can act as scaffolds, acting together with other factors into complexes that regulate many key nuclear activities. In mammals, alteration in BD function can lead to cell death or uncontrolled proliferation, and BD factors are often upregulated in some types of cancers and chronic inflammatory diseases, making them potential drug targets **[14].**

*T. cruzi* genome encodes eight bromodomain-containing proteins (BDF1–8), each with distinct features. When BDF1-3 KOs were attempted using CRISPR/Cas9 gene editing, it was only possible to obtain heterozygous mutants, which indicates that these proteins are essential at least for the epimastigote stage **[15]**. The expression of dominant-negative mutants provided additional evidence of the essentiality of these bromodomain-containing proteins **[15–17].** Among these eight *T. cruzi* BDs, BDF6 is notably divergent, with a unique 57-residue insertion in its AB loop and reduced similarity to canonical BD. Despite the BD and a monopartite Nuclear Localization Sequence (T_265_TRKGVT), *Tc*BDF6 does not possess any additional recognizable domain or feature.

We previously demonstrated that BDF6 is part of a TINTIN-like complex **[18]**. TINTIN is an autonomous module of the yeast histone acetyl transferase complex NuA4 (Nucleosome Acetyltransferase of H4), a key nuclear complex involved in numerous processes directly or indirectly associated with chromatin structure. Even though its function is not entirely understood, TINTIN is known to work as an independent sub-complex that can interact dynamically with canonical NuA4 and modulate its function. In yeast, it contains the only essential acetyltransferase **[18,19]**, and both NuA4 and TIP60, its homolog in mammals, have been implicated in transcriptional regulation, cell cycle progression and DNA repair, through histone acetylation **[19,20,21].**

Using a proximity labeling approach, we also identified structural orthologs of different chromatin remodeling complexes, including several NuA4 homologous components, clearly suggesting the presence of a NuA4-like complex in trypanosomatids (Rodriguez Araya, manuscript in preparation). In *T. brucei*, many of the same putative components of this complex were also identified by co-immunoprecipitation with BDF6 **[23]**. We have proved that BDF6 interact with two other proteins, we called MRGx (containing a MORF4-related gene domain) and MRGBP, to form a TINTIN-like complex **[18]**. By yeast two-hybrid assays, we demonstrated that the BC loop of BDF6 is involved in the interaction with MRGBP, which itself interacts with MRGx in a similar way to that in yeast and mammals these components do **[24]**. The evident structural conservation of *Tc*TINTIN complex contrasts with the low sequence similarity of BDF6, MRGx and MRGBP with their homologous from *Saccharomyces cerevisiae* (EAF5, EAF3, EAF7) or humans (BRD8, MRG15, MRGBP). This structural conservation clearly indicates that, even in the absence of evident sequence similarity, BDF6, Eaf5 and BRD8 could be considered functionally homologous factors, even though Eaf5 contains a pseudobromodomain rather than a canonical bromodomain **[18]**. In line with the functional importance of this protein family, the deletion mutant of Eaf7 exhibits strong growth deficiencies at 37 °C **[25]**, and BRD8 KO in mice leads to preweaning lethality, with its dysregulation associated with various cancers **[26]**.

Despite the increasing interest in epigenetic mechanisms in protozoan **[27]**, the functional roles of bromodomain-containing complexes in *T. cruzi* remain poorly characterized. A few previous studies have addressed essentiality in the epimastigote stage for some BDFs, however their contributions to vertebrate stages, particularly the intracellular amastigotes, remain unknown. In this work we obtained KO parasites for the gene *tcbdf6.* In the epimastigote stage these parasites can be cultured normally, even though they have a longer duplication time and a smaller size than Wild Type (WT) parental strain. By contrast, BDF6 deficiency heavily compromises both host-cell infection and amastigote development in cell cultures. Even though *TcBDF6 ^-/-^* parasites show normal *in vitro* metacyclogenesis efficiency, the metacyclic trypomastigotes (MTs) obtained present low infectivity and, once inside the cells, they differentiate into amastigotes within the parasitophorous vacuole without multiplying. This phenomenon is reversed by the episomal expression of the *tcbdf6* gene (add back strain). Finally, KO parasites show strong deficiencies to infect mice, an effect that is also mitigated in the add back strain.

Although functional studies on BDFs in *T. cruzi* are still limited, the unique phenotype of *TcBDF6* ^-/-^highlights its potential importance in the parasite’s biology and pathogenesis. Our findings also provide insights into the relevance of epigenetic mechanisms in parasite development and may lead to designing new therapeutic strategies targeting BD-containing proteins.

## Results

### BDF6 is not essential in the epimastigote stage, but its loss results in smaller parasites with slower duplication rates

By using CRISPR/Cas9-mediated genome editing and parasite cloning, we obtained a heterozygous (*TcBDF6* ^-/+^) and two homozygous (*TcBDF6* ^-/-^) KO clonal strains (Clones A and B). The knocking out strategy and PCR analysis of the clones are shown in supplementary figure 1. Since both complete KOs showed identical phenotypes after evaluation (see below), clone A was used to generate a complemented cell strain by reintroducing a version of the *tcbdf6* gene modified to avoid recognition by the CRISPR-Cas9 system (named thereafter *TcBDF6* ^-/-^pBDF6 or just “add back strain”).

To assess whether the absence of BDF6 affects *T. cruzi* epimastigote proliferation, we performed *in vitro* growth curve assays for five parasite strains: WT, heterozygous mutant (*TcBDF6* ^-/+^), two independent BDF6 KO clones (*TcBDF6* ^-/-^ Clone A and Clone B), and the add back strain (*TcBDF6* ^-/-^pBDF6). As can be observed in figure 1.A, the BDF6 KO clones displayed a consistently slower growth rate compared to WT and heterozygous parasites, with delayed entry into the exponential phase and reduced parasite densities at later time points. To quantify these differences, linear regression analyses were performed using data points from the exponential growth phase (48–120 hours), and duplication times were calculated. Both *TcBDF6* ^-/-^ clones exhibited significantly longer duplication times than the WT strain, and the heterozygous mutant showed an intermediate phenotype (Fig. 1.B). The add back strain partially restored the growth defect, achieving replication kinetics comparable to the heterozygous strain. In several attempts, we tested different concentrations of hygromycin in the culture, but we were unable to obtain the exact growth rate of the WT strain (not shown). This results are not surprising since the plasmid used to complement the add back strain, *p*TEX vector-derived gene expression, cannot be easily regulated **[29]**. To date, there are no reported plasmid-based systems in *T. cruzi* that allow tight, fine-tuned regulation of transgene expression levels, limiting the ability to precisely recapitulate endogenous protein abundance. On the other hand, it is also possible that overexpression of BDF6 may also have a deleterious effect on growth. All these findings demonstrate that BDF6 is not essential for growth but contributes to optimal epimastigotes proliferation under culture conditions.

**Figure 1.**
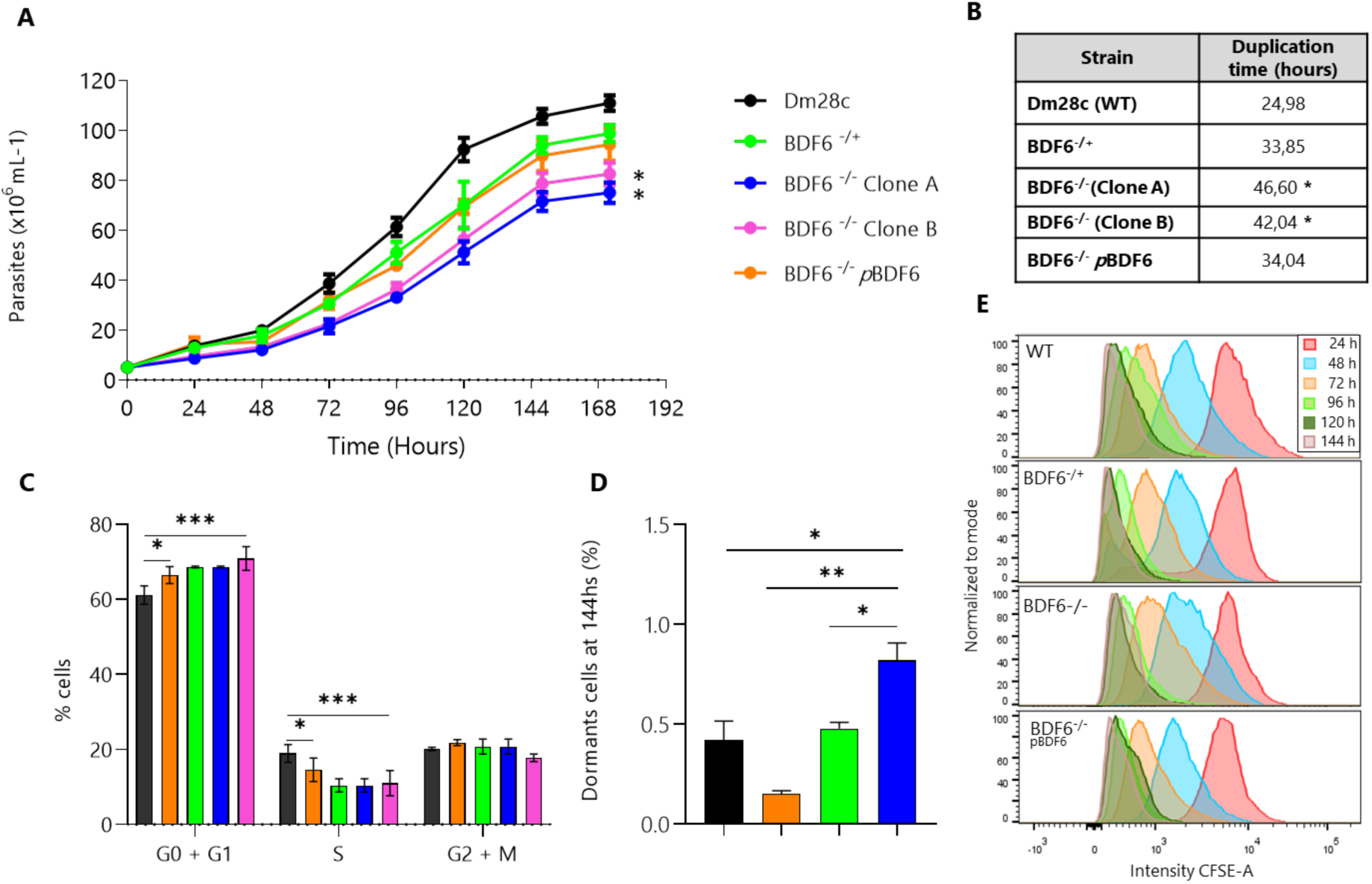
Deletion of *bdf6* impairs epimastigote replication and affects cell cycle progression. **A.** Growth curves and duplication times for different strains, displays a graph with parasite concentration (×10⁶/mL) and time in hours. Five curves are shown: WT Dm28c (black), BDF6^-/+^ (green), two independent BDF6^-/-^ clones (blue and magenta), and the complemented strain BDF6^-/-^ pBDF6 (orange). **B.** Table of duplication time: shows duplication times (in hours) for each strain **C.** Percentage of parasites in different stages on the cell cycle: bar plots show percentage of parasites in different stages on the cell cycle (G0/G1, S, and G2/M phases), measured by flow cytometry during exponential growth with IP straining. **D.** Percentage of dormants at 144 h: shows a bar graph quantifying dormant epimastigotes at 144 h for the same five strains. **E.** Histograms of CFSE fluorescent dye: it shows for each strain at 24, 48, 72, 96 and 144 h. Lower fluorescence indicates more cell divisions. In WT and the complemented strains, peaks shift progressively leftward over time, while KO clones retain a population with higher CFSE intensity at 144 h, indicating arrested or slowly dividing cells. Statistical significance is marked with asterisks (one-way ANOVA with Tukey’s test).

To investigate the potential cause of the reduced proliferation observed in *TcBDF6* ^-/-^ parasites, we measure DNA content in parasites in exponential phase of cultures as an estimation cell cycle of the parasites stage. As shown in figure 1.C, a higher proportion of *TcBDF6* ^-/-^ parasites (Clone A and Clone B) are accumulated in the G₀/G₁ phase compared to WT and heterozygous strains, suggesting a cell cycle delay at this checkpoint. The percentage of cells in S phase was lower in both KO clones, while the G₂/M populations remained comparable across all strains. Notably, the add back of BDF6 restored the G₀/G₁ and S phase distribution to levels like those observed in the heterozygous strain. These results suggest that loss of BDF6 causes a cell cycle delay at the G₁/S transition, restricting entry into S phase.

Given the reduced growth rate observed in *TcBDF6* ^-/-^ parasites, we investigated whether this phenotype could result from a higher proportion of non-dividing, dormant cells. To assess this, we performed a CFSE-based dormancy assay as described by **[30]**, evaluating epimastigotes from the four strains over a 144-hour period of culture. Briefly, CFSE-labeled parasites lose their fluorescence intensity with successive divisions, allowing to quantify the percentage of parasites that do not divide during culture. The BDF6 KO exhibited a significantly increased percentage of CFSE stained cells at 144 h compared to WT. The heterozygous *TcBDF6* ^-/+^ strain displayed an intermediate profile, consistent with a possible gene dosage effect (Fig. 1.D and 1.E). However, since the proportion of dormant cells remains in all strains below 1%, we considered that the subtle increase of dormant cells in *TcBDF6* ^-/-^strain is unlikely to be the main driver of the observed growth impairment. The observation that the add back strain showed fewer dormant parasites than the WT, but slower growth also reinforces the idea that the unregulated expression of BDF6 may itself have an effect on parasites.

To further evaluate the impact of the absence of BDF6 on epimastigotes, we performed ultra-expansion microscopy (U-ExM). As shown in Figure 2.A all strains of parasites exhibited a conserved morphology, with no evidence of granulation or structural abnormalities. However, KO epimastigotes appeared smaller than WT. When examined in detail, it was found that *TcBDF6* ^-/-^ epimastigotes were significantly shorter, but similar in width, than WT, while *TcBDF6* ^-/+^ and add back parasites exhibited an intermediate phenotype, with mean sizes falling between *TcBDF6* ^-/-^ and WT (Fig. S3). These size differences were also reflected in the light scattering values obtained from cytometry experiments designed to determine the cell cycle stages of the steady-state cultures (not shown).

**Figure 2.**
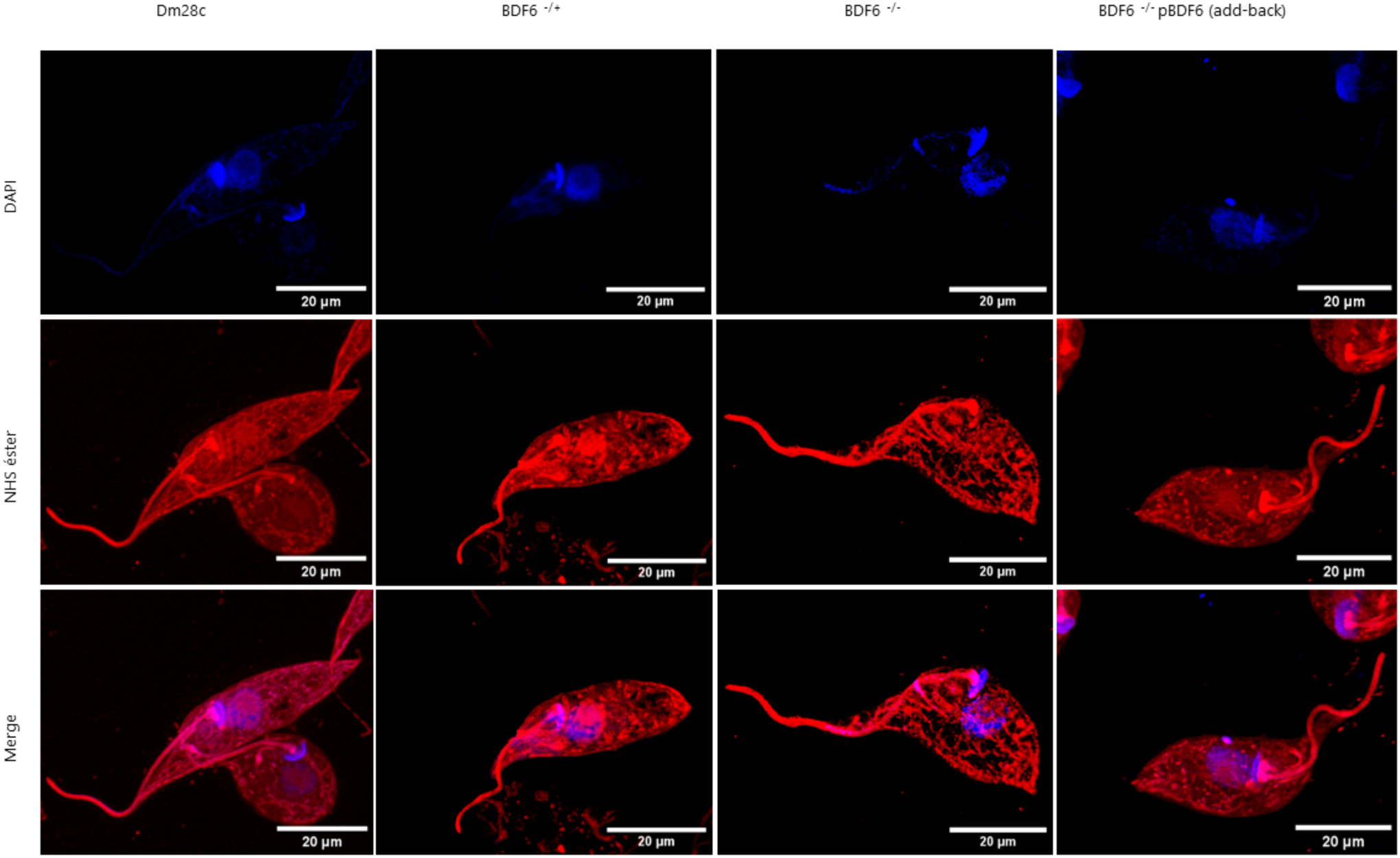
Ultra-Expansion Microscopy (U-ExM) of epimastigotes. Representative UExM images of WT Dm28c, heterozygous (*TcBDF6* ^-/+^), KO (*TcBDF6* ^-/-^), and add back (*TcBDF6* ^-/-^pBDF6) strains, stained with Alexa Fluor™ 647 NHS ester (red), which labels surface and accessible proteins, and DAPI (blue), which stains DNA. Scale bars: 20 µm

Then, in order to analyze the putative role of BDF6 on differentiation, we evaluated metacyclogenesis *in vitro*. All mutant strains produced metacyclic trypomastigotes (MTs) at levels comparable to WT Dm28c (Fig. S4) and the resulting MTs showed no morphological alterations under U-ExM (Fig. 3). Intermediate forms of metacyclogenesis, in addition to metacyclic trypomastigotes, were observed across all conditions as it was describe in Perdomo et.al 2025 **[31]**. Since these transitional forms appeared consistently in every case, they do not carry interpretative significance for the assessment of metacyclogenesis efficiency.

**Figure 3.**
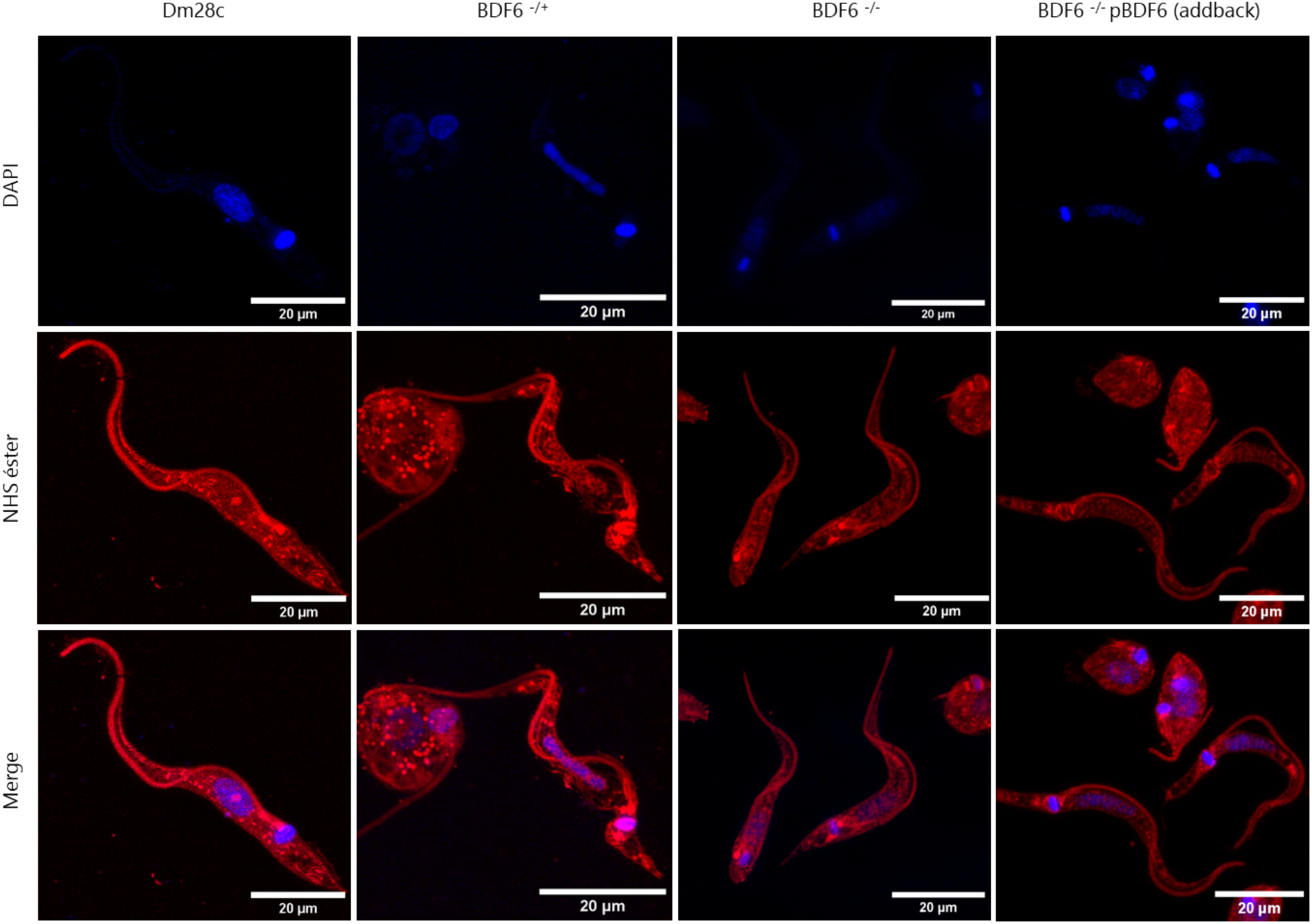
Ultra-Expansion Microscopy (U-ExM) of metacyclic trypomastigotes. Representative UExM images of WT Dm28c, heterozygous (*TcBDF6* ^-/+^), KO (*TcBDF6* ^-/-^), and add back (*TcBDF6* ^-/-^pBDF6) strains, stained with Alexa Fluor™ 647 NHS ester (red), which labels surface and accessible proteins, and DAPI (blue), which stains DNA. Scale bars: 20 µm

### BDF6 deficiency impairs the infectivity of trypomastigotes and compromises amastigote replication

Despite their conserved morphology, trypomastigotes from mutant strains were found to be functionally altered. As shown in figure 4.A and 4.B, the *in vitro* infection rate of the WT strain is in average 14% of the cells. In contrast, the infection rate for the heterozygous was reduced to around 2,5% and dropped to less than 1% for the homozygous mutant strain. Again, episomal expression of BDF6 in the add back strain restores the phenotype of the homozygous mutant, but inducing an increase in trypomastigotes infectivity comparable to that of the heterozygous mutant.

**Figure 4.**
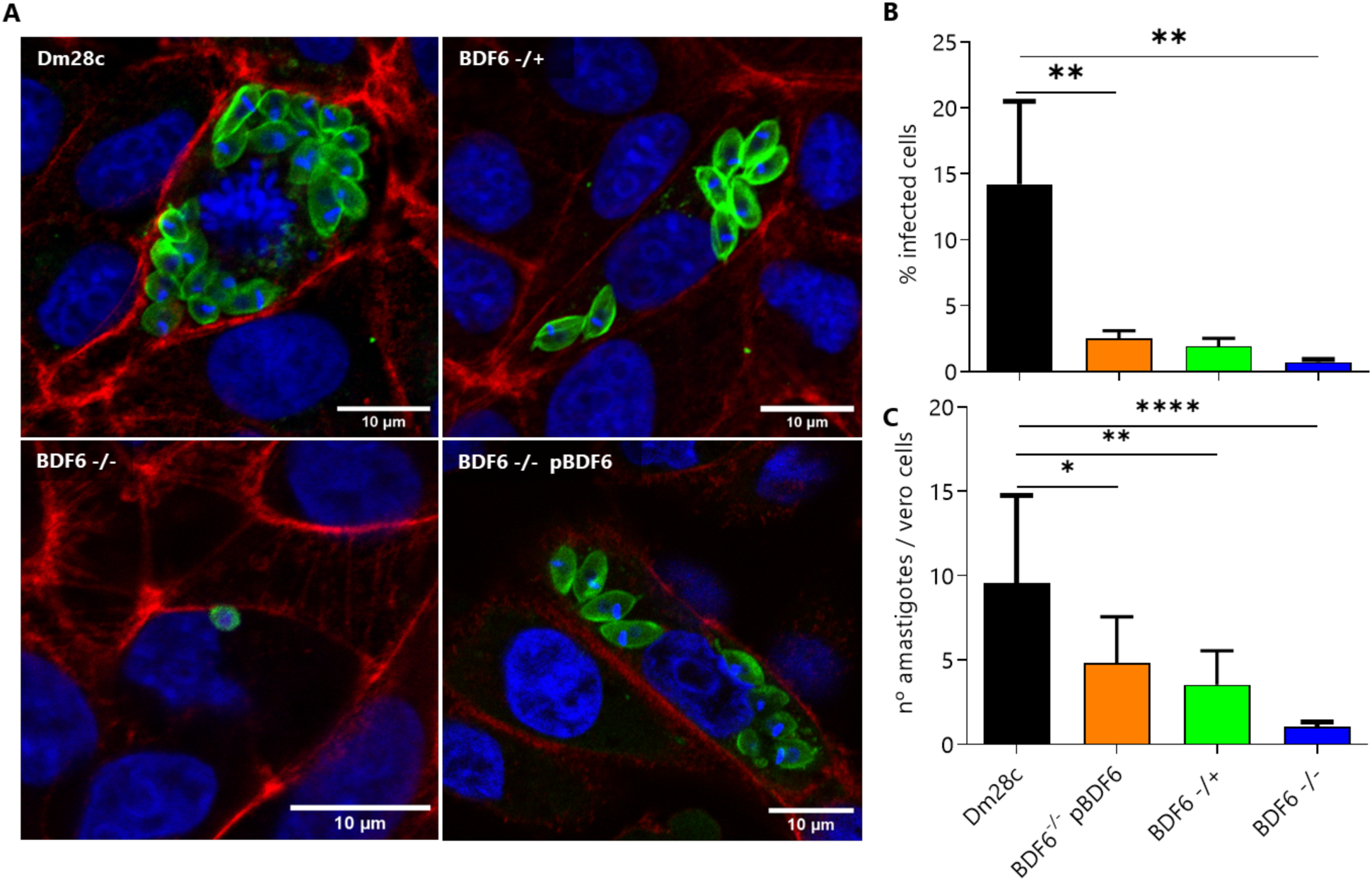
A. *In vitro* Vero cell infections. Representative confocal images of Vero cells infected with WT Dm28c, heterozygous (*TcBDF6* ^-/+^), KO (*TcBDF6* ^-/-^), and add back (*TcBDF6* ^-/-^pBDF6) strains. Intracellular amastigotes were visualized 72 hours post-infection (pi). Parasites were stained with anti-*T. cruzi* (green), host cell actin was labeled with phalloidin (red), and nuclei were counterstained with DAPI (blue). Scale bars = 10 µm (B) Quantification of infection rates (% of infected Vero cells) across the different strains. (C) Quantification of intracellular parasite load (number of amastigotes per infected Vero cell). Data represent mean ± SEM of three independent experiments. Statistical analysis was performed using one-way ANOVA followed by Tukey’s test (*p<0.05, **p<0.01, ***p<0.001, ****p<0.0001 compared to WT).

In a phosphoproteome analysis conducted along metacyclogenesis by Marchini et al. **[32]**, it was found that serine 97 of BDF6 is phosphorylated during this process (12 hours after parasite adhesion to the culture plate surface). To evaluate whether this post-translational modification could be related to BDF6-mediated trypomastigotes infection deficiency, we generated a new version of *bdf6* with a single replacement of serine 97 with alanine (*TcBDF6 ^-/-^* pBDF6_S97A_). However, episomal expression of BDF6_S97A_ showed no significant difference with the add back strain expressing BDF6, indicating that phosphorylation of this residue is not essential for trypomastigote infectivity (Fig. S2).

We also evaluated the number of intracellular amastigotes after *in vitro* infection of Vero cells. After 3 days of infection the WT strain displayed normal development of intracellular amastigotes. In contrast, in the *TcBDF6* ^-/-^ mutant strain we found only one amastigote per cell, while the heterozygous parasites showed an intermediate number of amastigotes per cell (Fig. 4.C). Interestingly, as well as observed for epimastigotes, *TcBDF6* ^-/-^ amastigotes seems to be also smaller than WT. Again, the add back strain showed intermediate number of the intracellular amastigotes, similar to that observed to the *TcBDF6* ^- /+^. Furthermore, the number of amastigotes of the *TcBDF6* ^-/-^ strain did not increase after several days’ post infection and no trypomastigotes were released after 11 days when cultured Vero cells began to lose viability (data not shown). These results highlight the importance of BDF6 not only for initiating infection but also for sustaining amastigote proliferation within host cells.

### After 48 hours of infection BDF6^-/-^ parasites remain within the parasitophorous vacuole

As can be shown in figure 4 and 5, amastigotes of the WT strain (green) replicates within host cells whereas amastigotes of the *TcBDF6* ^-/-^ strain remain within host cells but do not exhibit detectable replication. To investigate whether BDF6 KO parasites follow the classic transit of *T. cruzi* by escaping from the lysosomal-positive vacuole after host cell infection, we employed the lysosomal marker LAMP2 (Fig. 5). Notably, at 48 hours’ p.i., these *TcBDF6* ^-/-^ amastigotes still co-localize with LAMP2 (magenta), indicating their retention within the parasitophorous vacuole. These observations suggest that BDF6 is involved and may play a critical role in the intracellular development of *T. cruzi*, including the escape from the parasitophorous vacuole.

**Figure 5.**
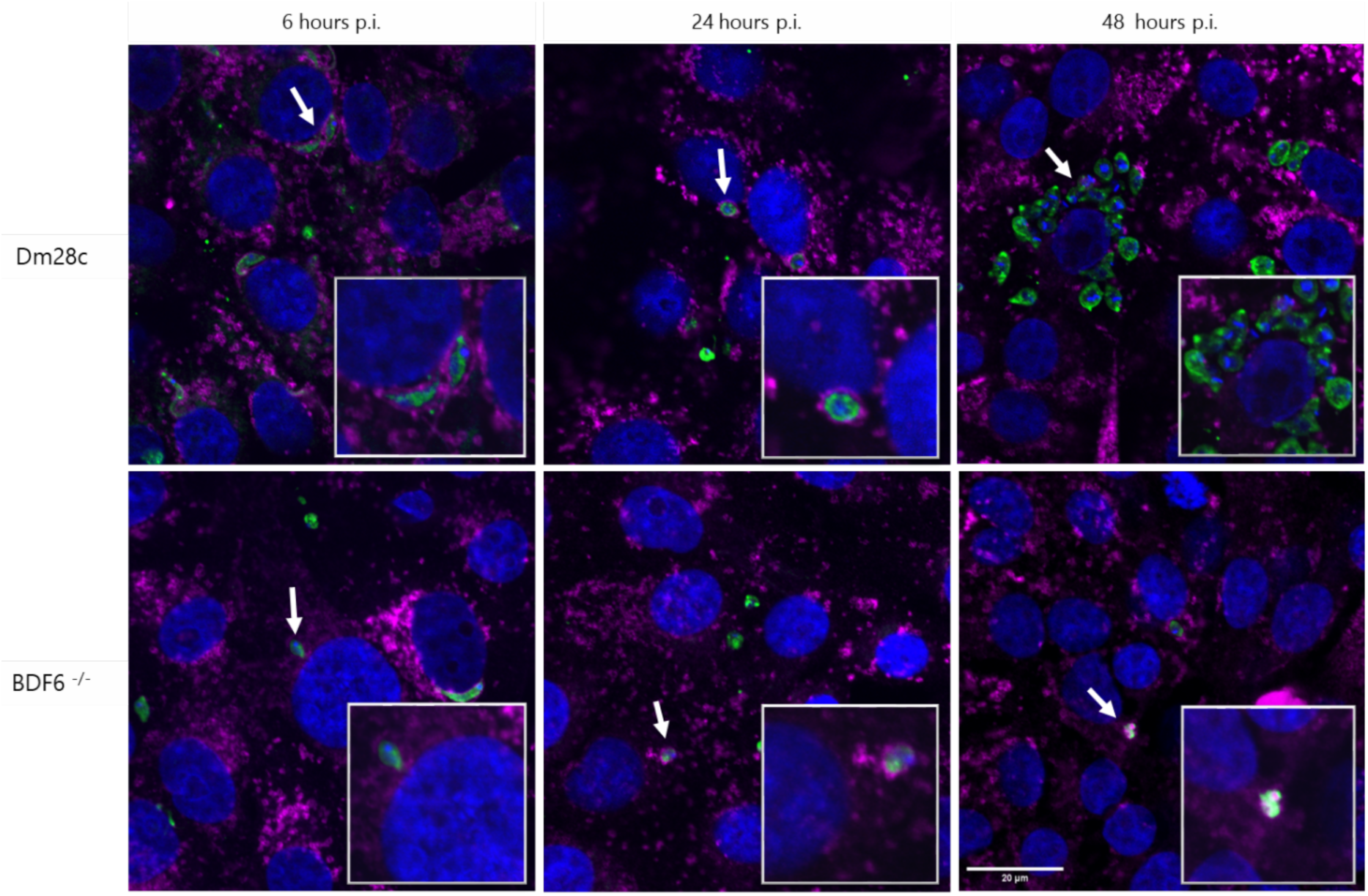
Immunofluorescence of infected Vero cell. Co-localization of lysosome marker and parasites were analyzed at 6, 24, and 48 hours’ post-infection. DAPI (blue), a-LAMP2 (magenta) and a-*T. cruzi* (green). Only one scale bar is shown (above right), as all the images have the same scale. Scale bar = 20um

One possible explanation for this observation is that *TcBDF6* ^-/-^ parasites could be retained at vacuole due to an increased susceptibility to oxidative stress generated by the host cell. To evaluate this, we exposed epimastigotes to different doses of H_2_O_2_ and nifurtimox as a source of superoxide radicals, and UV radiation. Under the conditions assayed, *TcBDF6* ^-/-^ parasites exhibited similar sensitivity to H_2_O_2_ and even displayed slightly greater resistance to nifurtimox compared to WT parasites (Fig. 6). These results indicate that BDF6 is not essential for resistance to oxidative stress, suggesting that the intracellular growth arrest observed is not due to an impaired ability to tolerate reactive oxygen species that could be produced by the host cell. Moreover, all strains showed similar sensitivity to UV irradiation, suggesting that the loss of BDF6 is not linked to defects in nucleotide excision repair mechanisms, one of the mechanisms proposed to trigger dormancy (see below). Instead, the observed amastigoteś growing defect probably results from an intrinsic inability to growth in the amastigote form rather than to a defective adaptation at the intracellular environment, resulting in retention in the parasitophorous vacuole.

**Figure 6.**
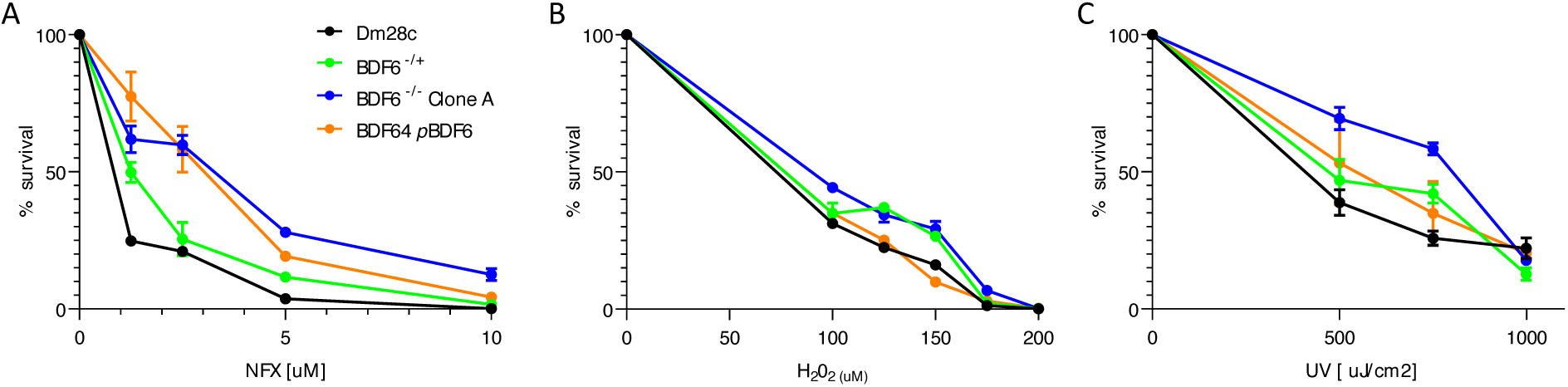
Sensitivity of epimastigotes to UV and oxidative stress. Epimastigotes of WT Dm28c, heterozygous (*TcBDF6* ^-/+^), KO (*TcBDF6* ^-/-^), and add back (*TcBDF6* ^-/-^pBDF6) strains were exposed to increasing doses of nifurtimox (A), H₂O₂ (B) and UV (C) and for 72 hours. Data represent mean ± SEM of four independent experiments.

### BDF6 is essential for parasite persistence *in vivo*

*In vivo* infection experiments performed in C57BL/6 and BALB/c nude mice revealed that BDF6 plays a critical role in infection dynamics. The WT strain used to generate the KO has low pathogenicity, and therefore, parasitemia is typically not observed in immunocompetent C57BL/6 mice up to day 30, before immunosuppression. However, after immunosuppression (starting at 31 d.p.i.), parasitemia rises at 50 d.p.i. in the WT strain (Fig. 7.A). In contrast, mice infected with both heterozygous (*TcBDF6* ^-/+^) and homozygous (*TcBDF6* ^−/−^) mutants, as well as the add back strain (*TcBDF6* ^−/−^pBDF6), failed to establish detectable parasitemia, indicating that BDF6 is necessary to sustain a microscopically detectable bloodstream infection, even under conditions of impaired immunity (Fig. 7.B). Spleen weight, used as a surrogate marker for immune activation, was significantly increased in WT-infected and immunosuppressed animals at day 53 p.i. (Fig. 7.C), consistent with a robust inflammatory response. In contrast, spleens from mice infected with *TcBDF6* ^−/−^, *TcBDF6* ^−/+^ and the add back strain remained similar in size to those of uninfected controls. Quantification of parasite burden in blood, heart and skeletal muscle tissues by qPCR (Fig. 7.B–E) at 53 d.p.i. showed that WT-infected mice harbored significantly higher parasite loads compared to *TcBDF6* ^−/−^ infected animals, where parasites were almost undetectable. The add back strain partially restored tissue colonization at levels similar to or even higher than the heterozygous strain. The apparent inconsistency of the results obtained with the add back strain with respect to what was previously observed may be due to a progressive loss of the plasmid after 53 days without the antibiotic selection pressure.

**Figure 7.**
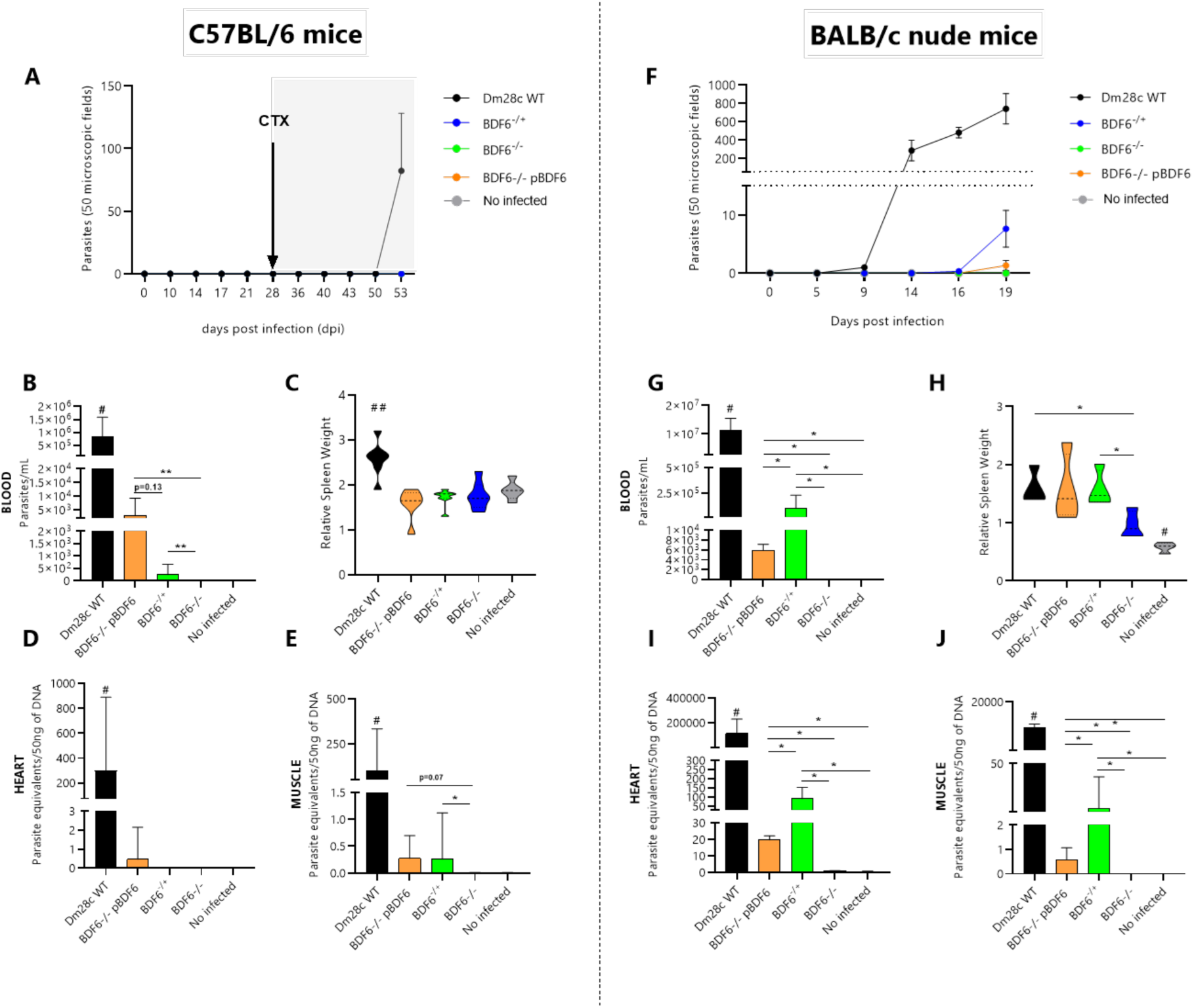
*In vivo* infections. (A–E) Infections in immunocompetent C57BL/6 mice subjected to immunosuppression with cyclophosphamide (CTX). (A) Parasitemia was assessed by direct blood examination at multiple time points post-infection. Only WT parasites exhibited detectable parasitemia following immunosuppression. (B) Quantification of parasite load in blood by qPCR revealed a significant reduction in *TcBDF6* ^-/-^ mutants compared to WT, while the add back strain (*TcBDF6* ^-/-^pBDF6) showed partial restoration. (C) Relative spleen weight, used as an indicator of immune activation, was significantly elevated in WT-infected mice but not in those infected with *TcBDF6* ^-/*-*^ mutants or the add back strain. (D, E) Parasite load in heart and muscle tissues, measured by qPCR, was markedly decreased in *TcBDF6* ^-/-^ infected mice, with partial recovery in the add back group. (F–J) Infections in immunodeficient BALB/c NUDE mice, which lack functional T cells. (F) Parasitemia kinetics showed robust parasite proliferation in WT-infected nude mice, while *TcBDF6* ^-/-^ mutants failed to establish significant parasitemia. Complementation partially restored parasitemia levels. (G) qPCR quantification in blood confirmed significantly reduced parasite levels in *TcBDF6* ^-/-^ infected BALB/c nude mice, with partial restoration in the add back strain. (H) Relative spleen weights remained low in *TcBDF6* ^-/-^ infected mice, comparable to uninfected controls, suggesting reduced immune activation. (I, J) Parasite burden in heart and muscle tissues was significantly reduced in *TcBDF6* ^-/-^ infected nude mice, with partial restoration upon complementation. Data are expressed as median ± range and were analyzed using non-parametric test (Kruskal-Wallis followed by Mann-Whitney U test). Results represent two independent experiments with 3–5 animals per group. Statistical comparisons between groups were performed using non-parametric tests. Significance levels are indicated as follows: *p < 0.05, **p < 0.01, ***p < 0.001 (between specified groups); #p < 0.05, ##p < 0.01 versus all other groups. Fig 7E and 7B show a marked difference in p-values. While one comparison reaches statistical significance (p < 0.05), the other shows a trend toward significance (p = 0.07), suggesting a potential biological effect despite not meeting conventional thresholds. Notably, the comparison appears different visually but is not statistically significant.

Despite differences in parasitemia levels, mice infected with WT, heterozygous, or add back strains and treated with cyclophosphamide showed a residual and comparable humoral response against *T. cruzi* antigens, as indicated by similar levels of specific IgG that persisted after 53 d.p.i. In contrast, mice infected with the homozygous KO strain exhibited IgG levels comparable to those of uninfected controls, which—as expected—did not develop a specific antibody response (Fig. S5).

To further assess the role of BDF6 in the absence of adaptive immunity, infections were also performed in nude mice. Nude mice lack mature T lymphocytes, resulting in a severely impaired adaptive immune response. While they retain functional B cells and innate immune components, their ability to establish an effective cell-mediated immunity is profoundly compromised. In this model, WT parasites reached high parasitemia levels starting around 9 d.p.i. (Fig. 7.F). *TcBDF6* ^−/−^ mutants again failed to establish parasitemia in nude mice, while those infected with the add back strain showed partial recovery, confirming the requirement of BDF6 for effective replication and systemic infection, even in immunodeficient hosts. Quantification of parasite burden in blood, heart, and skeletal muscle tissues by qPCR (Fig. 7. G, I–J) showed that WT-infected nude mice harbored significantly higher parasite loads compared to *TcBDF6* ^−/−^ infected animals, where the parasites were undetectable. The add back strain partially restored tissue colonization but, in this case, reaching parasite levels lower than those observed for the heterozygous strains. In nude mice, splenomegaly was evident after infection with WT, heterozygous and add back strains (Fig 7.H). In contrast, spleens from nude mice infected with *TcBDF6* ^−/−^ remained similar in weight to those of uninfected controls, indicating that *TcBDF6* ^−/−^ mutants consistently failed to establish an infection, even in the absence of T cell-mediated immune, highlighting the essential role of BDF6 for intracellular development.

## Discussion

In this study, we used the CRISPR/Cas9 genomic edition to investigate the impact of the bromodomain-containing protein BDF6 over *T. cruzi* life cycle. A common limitation of gene knockout approaches is the inability to generate viable epimastigotes if the target gene is essential across all life stages. In such cases, complete gene disruption may not be feasible. When homozygous KOs are obtained, the resulting phenotypes can result from severe to subtle growth defects, all along the life cycle. For instance, gene ablation studies have revealed defects in growth, differentiation, and infectivity in *T. cruzi* **[32,33]**. In other instances, heterozygous mutants, carrying only a single functional gene copy, exhibit significant phenotypes and are useful models to infer the gene’s function and significance. However, there are a few cases in which the phenotype is primarily manifested during the life cycle stages that occur in the mammalian host and shows a gene dosage-dependent phenotype. The disruption of active trans-sialidase genes, which does not affect the parasite’s ability to grow as epimastigotes or differentiate into metacyclic trypomastigotes, is one of them. These parasites can still invade host cells and escape from the parasitophorous vacuole, but exhibit impaired differentiation of amastigotes into trypomastigotes that do not egress from the host cell **[35]**. Another example is presented in this study. Deletion of the *tcbdf6* gene results in an exceptionally interesting and unique phenotype, unlike any previously described. BDF6 KO parasites proliferate as epimastigotes, with a subtle but clear difference in the size of the parasites, and differentiate normally into MT. However, these trypomastigotes are unable to infect efficiently and do not replicate as amastigotes once inside host cells, highlighting a life stage–specific essentiality not previously described for any BDF in this parasite. In parallel, the heterozygous*TcBDF6 ^-/+^* parasites show an intermediate phenotype between WT and KO parasites, and the add back cures the phenotype of the homozygous mutant strain into one similar to that of the heterozygous strain, supporting the conclusion that it is due to the lack of BDF6. As can be seen throughout this report, the restoration of the phenotype in the add back strain does not have the same intensity in all experimental situations. The expression of BDF6 in a non-regulated pTEX vector may explain this phenomenon. On the other hand, it is possible that increased expression of BDF6 also has deleterious effects on the parasites, as has already been observed for other BDFs **[28,36,36]**. Finally, in the mouse infection experiments, the lack of selection pressure does not allow for ensuring the persistence of the plasmid at a stable copy number, which explains why the complementation levels are less pronounced.

In trypanosomes, the regulatory mechanisms governing cell size and shape integrate the dynamics of the cytoskeleton, the cell cycle machinery and flagellar assembly. The microtubule-based cytoskeleton establishes a scaffold that supports cell shape, directs organelle positioning and facilitates the asymmetric partitioning of cellular contents during division **[37,38]**. In parallel, the flagellum influence cell polarity and modulates overall cell dimensions **[4,39,40]**. Regulation of cell size through genetic pathways provides an additional layer of control that intersects with the physical dimensions imposed by cytoskeletal and flagellar structures **[40,41]**.

Campbell and de Graffenried recently demonstrated that the division of *T. cruzi* is asymmetric, generating a parasite of shorter length (but the same width) that contains a new flagellum, and a parasite similar in size to the epimastigotes in a state of division, which contains the flagellum from the original cell **[42]**. This size difference is due to the smaller size of the new flagellum. Following cytokinesis, the size of the resulting epimastigotes equalizes through a process of parasite elongation, accompanied by an elongation of the new flagellum. Unlike many eukaryotes, trypanosomes cellular growth during the G1 phase is just restricted to this elongation of the new flagellum derived parasites **[40, 37]**, with main parasite growth occurring during the S and G2/M phases **[43,40,41]**. The fact that lack of BDF6 generates smaller epimastigotes, without altering their morphology as well as normal MT, indicates that there are no deficiencies at the level of the cell body cytoskeleton but have shorter flagella. Since parasites are shorter, but similar in width make us to assume that subpellicular corset is conveniently conserved and suggests some kind of alteration during the cell cycle evolution that govern flagellum elongation. However, even though the accumulation of parasites at G0/G1 can explain the extension of the duplication time, there is not a clear hypothesis to explain size decrease of the parasites.

A possible interpretation of the *TcBDF6 ^-/-^* parasites phenotype could be referred to their higher predisposition to be in a dormant state. Dormancy is defined as the ability to enter a quiescent state with reduced metabolic activity, more resistant to stressful environmental conditions, such as drug treatment, resulting in the persistence of the infection [46,47]. Although this phenomenon was first described in intracellular amastigotes, where it is relevant for infection persistence, it has also been observed in epimastigotes **[48]**. The transition into and out of dormancy in *T. cruzi* is thought to be a spontaneous and stochastic process, and the mechanisms driving dormancy remain unknown **[49]**. The only established mechanistic hypothesis until now suggests that spontaneous DNA damage activates DNA repair mechanisms, leading to cell cycle arrest until the repair is complete [30,48]. However, the dormancy hypothesis cannot explain the *TcBDF6* ^-/-^ phenotype. For one side, while a higher number of dormant epimastigotes was observed in KO, its low amount doesńt explain mathematically the increased average doubling time. Moreover, in the same experiment we observed that add back strain has lower dormant parasites that the WT, but still growth slowly. On the other side, the observation that *TcBDF6* ^-/-^ parasites do not show increased susceptibility to reactive oxygen species or UV irradiation compared to the parental strain, which also argues against the hypothesis of amastigotes being in a dormant state, at least not triggered by DNA damage. Finally, it should be stated the dormant hypothesis cannot be tested in *TcBDF6* ^-/-^ amastigotes, because their replication is completely impaired (Fig 4A). Moreover, amastigotes do not appear to “awaken” from the putative dormant state after 11 days in Vero cells, nor after more than 20 days in immunosuppressed nude mice.

Although direct evidence linking epigenetic mechanisms, cell-cycle control, and quiescence in trypanosomatids remains unclear, emerging studies offer intriguing clues. For instance, ATAC-seq in *Leishmania donovani* has revealed stage-specific chromatin accessibility, with euchromatin predominating in fast-growing promastigotes and heterochromatin in slowly proliferating amastigotes **[50]**. In *Trypanosoma brucei*, acetylation of H4K4 is dynamically regulated during the cell cycle by HAT3 **[51]**, while in *T. cruzi*, mass-spectrometry studies have shown increases in acetylation at H4 N-terminal lysines (K2, K5, K10) during S phase **[52]**. These observations, together with the fact that BDF6 is taking part of a NuA4-like acetylase complex, support the hypothesis that epigenetic regulation might intersect with pathways controlling cell-cycle progression and entry into low-proliferative or quiescent states in trypanosomatids. Although the precise cause of the observed phenotype lies beyond the scope of this study and requires further investigation, we speculate that the effects of BDF6 absence on MT infectivity and amastigote proliferation may arise from distinct mechanisms. Metabolic or genetic defects affecting critical pathways are unlikely, as such alterations would have severely impacted epimastigotes growth. Furthermore, the differentiation from MT to amastigotes appears to proceed normally. Infectivity and escape from the endocytic vacuole, processes that are largely driven by surface proteins, might be selectively impaired in the absence of BDF6 by unknown mechanisms. The arrest in intracellular amastigote replication, which does not result from increased sensitivity to oxidative stress, can instead reflect defects in stage-specific cell cycle control.

Finally, since the slower growth in epimastigotes and the completely lack of cell division in amastigotes can be both associated to alterations in the cell cycle, it should be considered the possibility that these phenotypes even completely different in intensity, may be related. As already mentioned, cell cycle regulation in *T. cruzi*, entails significant differences compared to other eukaryotes. A number of key molecular components such as cyclins and cyclin-dependent kinases (CDKs) have been identified in this parasite. However, transcriptomic analysis along the cell cycle, indicates that trypanosome cell cycle regulation relies on a complex regulatory network involving phosphorylation and diverse and overlapping roles for cyclins and kinases, unlike other eukaryotes where CDK activity is often regulated by cyclins in a more linear fashion **[53–55]** This regulatory network besides sharing several proteins, appears to be different in epimastigotes and amastigotes. In this context, it can be hypothesized that the lack of BDF6 could be affecting this network in such a way that it generates a much more pronounced effect in amastigotes than in epimastigotes. For example, it has been shown that the expression of non-acetylable versions of H4 at K10 or K14 induces, in epimastigotes, decreased S-phase progression resulting in more cells in G1 **[56].** Considering that BDF6 seems to be part of a NuA4-like acetylase complex, responsible for H4 acetylation in other cell types, we could infer that a similar type of phenomenon could be occurring but affecting amastigotes to a greater degree than epimastigotes.

While the underlying mechanisms remain to be elucidated, our findings support the idea that BDF6 is implicated in epigenetic processes that are critical for successful amastigote development via its participation in the TINTIN/Nua4 complexes. The findings presented here open promising avenues for future research with potential implications for Chagas disease treatment and prevention. Previously, we demonstrated that inhibitors of BET family BDs exhibit trypanocidal activity. Among the eight *T. cruzi* BDFs, BDF1 and BDF3 were classified within this family. To date, BDF6 has not been assigned to any known BD family. However, its involvement in an evolutionarily conserved complex related to NuA4/TIP60 suggests a possible functional relationship with BRD8, which together with BAZ1B, BRWD1, BRWD3, CREBBP, EP300, PHIP, and WDR9, belongs to subgroup III of human bromodomain-containing proteins **[57]**. The development of specific inhibitors targeting BRD8 is relatively recent and less advanced than BET family inhibitors (group II), and their efficacy against *T. cruzi* has not yet been explored. It is noteworthy that since BDF6 is essential for intracellular amastigote development, it is an ideal point of intervention for designing new trypanocide therapies. Additionally, a highly attenuated *T. cruzi* strain, like *TcBDF6* ^-/-^, that grows efficiently as epimastigotes and differentiates into trypomastigotes, yet fails to maintain infection, could serve as a potential vaccine candidate for immunizing domestic animals involved in the maintenance of the domiciliary and peridomiciliary transmission cycles of *T. cruzi*.

## Materials and methods

### Parasite cultures

*T. cruzi* Dm28c WT epimastigotes were cultured at 28°C in complete LIT medium (5g/L liver infusion, 5g/L bacto-tryptose, 68mM NaCl, 5.3mM KCl, 22 mM Na_2_HPO4, 0.2% (w/v) glucose, and 0.002% (w/v) hemin) supplemented with 10% (v/v) fetal calf serum (FCS) (Internegocios S.A., Argentina)].

### *Tc*BDF6 KO strains generation

To knockout the *tcbdf6* gene, CRISPR/Cas9-mediated genome editing was done, as previously described **[58,59]**. Briefly, the Cas9/pTREX-n vector (Addgene, catalog number: 68708) was used to clone a specific sgRNA sequence to target *tcbdf6* gene (TriTrypDB ID: *Tc*C4B63_41g80), which encodes the BDF6 protein. The sgRNA carrying the specific protospacer sequence was amplified from the pUC_sgRNA plasmid using the oligonucleotides sgRNAFw and sgRNARv (Table S1) and cloned into Cas9/pTREX-n plasmid. Donor DNA was generated by PCR using the blasticidin-S deaminase gene (*Bsd*) as template. Ultramers used were *TcBDF6*_BSD_KO and *TcBDF6*_BSD_KO Reverse (Table S1) and included the homology region for DNA recombination. *T. cruzi* WT epimastigotes were co-transfected with donor DNA and Cas9/pTREX-n plasmid, and mutant parasites (*TcBDF6* ^-/-^) were selected by adding G418 (250 µg/mL) and Blasticidin (5 µg/mL) to the culture medium, ensuring continuous CRISPR-Cas9 system operation.

Subsequently, limiting dilution cloning assays were performed. Mutant genotypes were confirmed by PCR using Val_BDF6_KO_Fw and Val_BDF6_KO_Rv oligonucleotides (Table S1). PCR product for WT strains showed one band of 1391 bp, while homozygous (*TcBDF6* ^-/-^) mutant strains displayed one band of 1791 bp, and heterozygous (*TcBDF6* ^-/+^) strains showed both bands. This result was interpreted as a successful homologous recombination of the *Blasticidin* resistance gene into the *tcbdf6* gene locus. Three clones were selected, one heterozygous (*TcBDF6* ^-/+^), and two homozygous, named *TcBDF6* ^-/-^ Clone A and *TcBDF6* ^-/-^ Clone B.

### Complementation of KO strains

*T. cruzi* genomic DNA (gDNA), extracted from epimastigotes WT Dm28c, was used for the amplification of *tcbdf6* gene. PCR amplification employed a forward oligonucleotide TcBDF6_MutCas9 (Table S1) designed to introduce silent mutations in the Cas9 recognition site at the 5′ region of the *tcbdf6* gene, by modifying the third bases of the first six codons and introducing a *Bam*HI restriction site upstream of the gene. These mutations preserve the amino acid sequence but prevent recognition by the CRISPR-Cas9 system in KO strains. The reverse oligonucleotide TcBDF6_*Eco*RV (Table S1) bound to the 3′ regions of the gene and introduced an *Eco*RV restriction site. The PCR products and the pENTR3C vector were digested with *Bam*HI and *Eco*RV restriction enzymes, resolved on a 1% agarose gel, and purified using the Wizard® SV Gel and PCR Clean-Up System (Promega). The purified fragments were ligated and transformed into *E. coli* DH5α cells. Plasmid pENTR3C-BDF6-mc9 was isolated.

We transferred from the entry vector (*pENTR3C*-BDF6-mc9) into the destination vector *pTEX-GW*, shearing Hygromycin resistance by LR recombination reaction using Clonase® enzyme mix (Invitrogen). *pTEX-GW* was linearized with *Eco*RI (Promega) for 4 hours at 37°C to enhance recombination efficiency. The resulting construct was transformed into *E. coli*, purified, and subsequently used for transfection into *T. cruzi* to complement mutant strains and study BDF6 function.

### Growth curves

An initial inoculum of 5x10^6^ epimastigotes/ml was cultured in LIT medium supplemented with 10% FBS, 0.8% glucose, and 0.002% hemin. Daily, aliquots of the culture were taken, and diluted in a 4% PBS-formaldehyde solution. Parasites were counted by Neubauer chamber or hematological counter (Wiener Lab Counter 19) in the white cells channel.

### Dormancy assay

Dormancy assays were performed as described by **[30]**. Briefly, epimastigotes in the exponential growth phase were harvested and washed twice with 1X PBS. Parasites were suspended in 1 mL of PBS containing 1 µL of CFSE (CellTrace CFSE, Thermo Fisher) stock solution. After 20 minutes of incubation in the dark at 28°C, the reaction was stopped adding 200 µL of FBS, followed by centrifugation at 3500 × g for 10 minutes. Parasites were resuspended in complete LIT medium at a final concentration of 5 × 10⁶ parasites/mL. To assess fluorescence intensity at different time points, samples of approximately 1 × 10⁶ parasites were collected daily from day 1 (24 hours) till day 6 (144 hours). Samples were centrifuged, washed with 1X PBS, and fixed for 15 minutes with 50 µL of 4% filtered paraformaldehyde (PFA), centrifuged, and suspended in 1X PBS for analysis.

Ten thousand events per sample were acquired using BD FACSAriaII (BD Biosciences). The percentage of dormant epimastigotes was calculated by counting the burden of parasites after 144 hs retaining maximum fluorescence intensity, which was previously determined by the intensity of fluorescence in parasites at 24 hours. The results were analyzed using FlowJo software (Bd Biosciences).

### Quantification of the size of epimastigotes

Fixed epimastigotes from exponential stage were mounted on microscope slides and stained with Giemsa solution, in a dilution 10% (v/v), for 20 minutes. Bright-field microscopy images were captured at 40× magnification and saved in TIFF format to preserve quality for analysis. Image analysis was performed using the ImageJ software (version 1.54p). Images were converted to grayscale (Image > Type > 8-bit) to facilitate segmentation. A threshold was applied (Image > Adjust > Threshold) to clearly distinguish epimastigotes from the background. The threshold settings were adjusted to ensure precise delineation of the parasite boundaries. Subsequently, the areas corresponding to epimastigotes were selected using the Analyze Particles function (Analyze > Analyze Particles). Parameters for particle size and circularity were set to filter out noise and exclude non-epimastigote structures. Only objects matching the expected morphology and size of epimastigotes were included in the analysis. For each selected epimastigote, measurements of individual cell areas were generated by ImageJ (version 1.54p).

### Ultra resolution-expansion microscopy

To visualize morphological characteristics of epimastigotes and metacyclic trypomastigotes, the methodology described by **[60]** was used. Briefly, one million parasites were washed with PBS, settled on poly-D-lysine-coated coverslips, and fixed with 4% formaldehyde and acrylamide. Gelation was performed using a monomer solution, followed by polymerization on ice and incubation at 37 °C. The gels were detached using a denaturation buffer and then incubated at 95 °C for 1.5 hours. After expansion in Milli-Q water overnight, gels were washed in PBS, stained with DAPI/NHS ester Alexa Fluor 647 (Thermo Fisher Scientific, Carlsbad, CA, USA) overnight at 4 °C, and subjected to a second round of expansion with multiple water exchanges. Images were acquired with a confocal Zeiss LSM880 microscope. ImageJ software was used to pseudocoloring and processing all images **[61]**.

### Cell cycle analysis of epimastigotes by flow cytometry

To determine cell cycle distribution in *T. cruzi* epimastigotes, we performed flow cytometric analysis based on DNA content using propidium iodide (PI) staining. Briefly, epimastigotes from exponential phase cultures were collected, washed twice with PBS, and fixed in cold 70% ethanol and incubated on ice for 40 minutes to ensure optimal fixation and permeabilization. Cells were centrifuged at 3,000 × g for 10 minutes and washed twice with PBS to remove residual ethanol. The pellet was then resuspended in PI staining solution (50 µg/mL PI, 100 µg/mL RNase A, and 0.1% Triton X-100 in PBS). Samples were incubated for 30 minutes at 37 °C in the dark. Stained cells were analyzed on a BD FACSAriaII cytometer (BD Biosciences) equipped with a 488 nm laser. PI fluorescence was collected in the FL2 channel (585/42 nm filter). A total of 10,000 events per sample were recorded and cell cycle distribution was determined using FlowJo software (Version 10). Debris and aggregates were excluded by appropriate gating, and the percentage of cells in G₀/G₁, S, and G₂/M phases was calculated based on DNA content histograms.

### Oxidative and UV stress assays

To evaluate the response of epimastigotes to oxidative stress, parasites in exponential growth phase were collected (5 × 10⁶ parasites/mL), centrifuged at 4,000 × g for 10 minutes, and washed twice with 1× PBS to eliminate residual culture medium. For hydrogen peroxide (H₂O₂) treatment, parasites were incubated for 30 minutes in PBS containing 0, 50, 100, or 150 µM H₂O₂, then washed, suspended in LIT medium (5 × 10⁶ parasites/mL), and plated in triplicate (500 µL/well) in 24-well plates. For drug-induced oxidative stress, 500 µL of epimastigotes (10×10⁶ parasites/mL) from WT, *TcBDF6* ^-/+^, and *TcBDF6* ^−/−^ strains were seeded in triplicate and treated for 72 hours at 28 °C with increasing concentrations of benznidazole or nifurtimox. For UV-induced stress, parasites were exposed to increasing doses of UV radiation (500, 750, and 1000 µJ/cm²), then resuspended in LIT medium and incubated for 72 hours. After treatment, parasites were fixed with 3.7% formaldehyde and counted using a Neubauer chamber to determine % survival.

### *In vitro* metacyclogenesis

Metacyclogenesis of all strains was evaluated as described by **[31]**. Briefly, cultures in exponential phase with an initial concentration of 5x10^6^ epimastigotes/ml, parasites were incubated at 28°C in 24-well plates, in quadruplicate, with 500 µl of LIT medium (0.8 % glucose) at pH 6 for 7 days. Subsequently, aliquots of the culture were taken and diluted in a 4% PBS-formaldehyde solution, and the different parasite stages were counted in a Neubauer chamber. To determine the percentage of metacyclogenesis, the number of trypomastigotes was counted and expressed as a proportion of the total number of parasites.

### *In vitro* infection of Vero cells

Vero cell line (ATCC CCL-81) was cultured in DMEM (Dulbecco’s Modified Eagle’s Medium powder, Gibco™, Thermo Fisher Scientific, Cat. No. 12100-046), supplemented with 10% (v/v) heat-inactivated FBS, at 5% CO_2_, at 37°C. Metacyclic trypomastigotes (MT) were obtained as previously mentioned and quantified. Then, in 24-well plates containing 12-mm round glass coverslips, 20,000 Vero cells were seeded per well in DMEM supplemented with 2% FBS and incubated for 24 hours at 37°C, 5% CO₂. Cells were infected with MT at a multiplicity of infection (MOI) of 10:1. After 16 hours of incubation (ON), non-adherent parasites were removed by washing twice with PBS 1X, and the DMEM medium was replaced. Cultures were maintained by five days, infected cells were washed twice with PBS and fixed with PFA 4%. Parasites were visualized by Giemsa or DAPI staining and immunostaining using mouse *a-T. cruzi* antibody. Images were acquired with a Zeiss LSM 880 confocal microscope. Microphotographs were taken, and the percentage of infected cells and the mean number of intracellular amastigotes per infected cell were quantified. Three hundred cells from each quadruplet were evaluated, and the experiment was repeated three times.

### Colocalization of amastigotes and lysosomal markers

Colocalization assay was performed using antibodies against the endogenous lysosomal protein LAMP2 and a mouse *a-T. cruzi* antibody **[62]**. Infections were performed as previously described. For lysosomal detection, cells were incubated overnight at 4°C with a primary anti-LAMP2 antibody diluted in 1% PBS-BSA. Samples were washed three times with PBS and subsequently incubated for 1 hour at room temperature, protected from light, and stained with an Alexa 555-conjugated secondary antibody. A second immunostaining step was performed using a primary anti-*T. cruzi* antibody followed by a FITC-conjugated secondary antibody to detect amastigotes. After three final PBS washes, samples were mounted with a DAPI-containing medium to stain the nuclei of both Vero cells and amastigotes **[63,64]**. Coverslips were placed on the samples and sealed. Images were acquired by confocal microscopy. Colocalization of LAMP2 (lysosomes) and anti-*T. cruzi* signals were analyzed to evaluate the retention of amastigotes within the parasitophorous vacuole.

### Mice infections

The infection experiments were performed on C57BL/6 male mice (6–8 weeks old) and on Balb/c male nude mice (6–8 weeks old, immunodeficient, lacking T cells). Animals were housed in HEPA-ventilated racks with *ad libitum* access to food and water. All procedures were conducted by institutional and national ethical guidelines for animal care and use. The protocol was approved by the ethical committee for animal experiments (CICUAL) of the Facultad de Ciencias Médicas at the Universidad Nacional de Rosario (Res N° 911/2023).

Mice were divided into five experimental groups (n=3-6/group), each named according to the *T. cruzi* strain used for infection: Uninfected control (negative control), WT strain (positive control strain), *TcBDF6* ^-/+^ (heterozygotes strain), *TcBDF6* ^-/-^ (KO strain) and *TcBDF6* ^-/-^pBDF6 (add back strain). For the infection, MT were obtained as described above. One million MT per mouse were injected intraperitoneally in PBS+Glucose (100 mg/dL). Control group received the same amount of vehicle. The experiment was conducted over 53 days for C57BL/6 mice and 21 days for Balb/c nude mice, monitoring parasitemia regularly post-infection (pi).

### Blood parasite burden

Parasitemia was assessed by quantifying circulating trypomastigotes in peripheral blood samples obtained via tail vein puncture using heparinized tips. Fresh blood samples were mounted directly onto microscope slides without fixation or staining and examined immediately under a light microscope. Trypomastigotes were counted in a defined volume of blood to estimate parasite burden as previously reported **[65]**.

### C57BL/6 mice immunosuppression

WT, *TcBDF6* ^-/+^, *TcBDF6* ^-/-^ and *TcBDF6* ^-/-^pBDF6 infected C57BL/6 mice (n=3-6/group) were treated with three doses of cyclophosphamide (200 mg/kg, IP), administered every four days, starting on day 31 post-infection, to induce immunosuppression during the course of infection. An uninfected group of mice (n=4) was subject to the same procedure.

### Detection of parasite burden in tissues

Hearts and skeletal muscles were collected and stored at -80°C for subsequent DNA extraction, following the method previously described **[66]**. Briefly, total DNA was extracted from each sample after tissue disaggregation with a lysis buffer (10 mM Tris-HCl; 0.1 M NaCl; 10 Mm EDTA; 0.5% SDS, 300 µg/mL proteinase K from Inbio Highway®. pH = 7.6). The samples were then heated for 2 h at 55 °C, and extracted twice with a phenol–chloroform–isoamyl alcohol mixture (25:24:1) (Sigma Chemical Co., St. Louis, MO, USA). Cold ethanol (AAPER Alcohol and Chemical Co., Shelbyville, KY, USA), twice the volume of the extracted sample, was then added and the samples were placed at −80 °C for ON. Samples were centrifuged for 30 min at 8000g and washed with 70% ethanol, vacuum dried, and then resuspended in 100 µL of sterile miliQ water. DNA samples were adjusted to a final concentration of 25 ng/µL and were used as a template for the qPCR reactions using the specific oligonucleotides for *T. cruzi*: TCZ-F and TCZ-R (Table S1), which recognize a satellite sequence of 182 bp present in the parasite genome. To generate the *T. cruzi* quantification curve, 25 mg of uninfected mouse hearts or muscles were mixed with 1x10^6^ trypomastigotes. Total DNA was extracted, and the final concentration was adjusted to 25 ng/ml. Following, the standard curves were constructed by serial dilutions using DNA from uninfected tissues (25 ng/ml) as diluent. PCR reactions were performed using a HOT FIREPol EvaGreen qPCR Mix Plus kit (Solis Biodyne®) with a StepOneTM Real-Time PCR Systems (Applied Biosystems®) instrument. The PCR conditions consisted of an initial denaturation at 95 °C for 10 min, followed by 40 cycles of 95 °C for 10 s, 50 °C for 15 s, and 72°C for 15 s, with fluorescence acquisition at 81°C for 15 s. Amplification was immediately followed by a melt program with an initial denaturation of 10 seconds at 95°C, 60°C for 20 seconds, and then a stepwise temperature increase of 1 °C/s to 90°C. The parasite load was expressed as equivalent parasites in 50 ng of murine DNA.

### qPCR quantification of parasite load in blood

Blood samples were immediately mixed with two volumes of lysis buffer containing 6M guanidine hydrochloride (Sigma®) and 200 mM EDTA, pH 8.0 (GE buffer). A pGEM-T plasmid (Promega®) was used as a heterologous internal standard (IS) and all samples were mixed before DNA purification with 1 pg of the IS. DNA was purified with High Pure PCR Template Preparation Kit (Roche®) following the manufacturer’s instructions. PCR was performed as already mentioned. A quantification curve for *T. cruzi* was generated by spiking uninfected mouse blood with 1 × 10⁶ Tulahuen strain trypomastigotes or 1000 pg of IS DNA, followed by immediate processing as described. Total DNA was extracted, and both standard curves were constructed by serial dilutions using uninfected mouse blood DNA as diluent. The parasite load was expressed as number of *T. cruzi*/ml of blood as described in **[67].**

### ELISA for anti-*T. cruzi* antibody detection

Blood samples were collected at day 53 p.i. by cardiac-punction with anticoagulant and plasma was obtained by centrifuging at 5000 rpm and stored at −20°C until analysis. ELISA microplates coated with *T. cruzi* lysate (Wiener Lab®) were used. Plasma samples were diluted 1:32 in 0.5% BSA and incubated in duplicate for 1 hour at 37°C. Specific anti-*T. cruzi* antibodies were detected using a peroxidase-conjugated goat anti-mouse IgG antibody (diluted 1:2000; Sigma®), incubated for 1 hour at 37°C. After washing, TMB substrate was added, and the reaction was stopped with sulfuric acid. Absorbance was measured at 450 nm with correction at 545 nm in an ELISA reader (Epoch Biotek Instruments®).

### Statistical analysis

Statistical analysis was performed with GraphPad Prism 3.0 software (GraphPad Software, La Jolla, CA, USA). For *in vitro* experiments, comparisons were made using One-Way Analysis of Variance (ANOVA) followed by Tukey’s post hoc test. For *in vivo* experiments in mice, non-parametric tests were used: Kruskal-Wallis followed by Mann-Whitney U test. Depending on the variable analyzed, data are expressed as mean ± standard error of the mean (SEM) or median ± range. Results represent one of three independent experiments performed on different days using different culture batches. At least 3 biological replicates per experimental group were used in each experimental round. Significance was set at p<0.05.

## Supporting information

Table S1, Fig. S1, Fig S2, Fig S3, FingS4, Fig S5

## Acknowledgments

We would like to thank Rodrigo Vena for technical confocal microscope assistance, Dolores Campos for Vero Cells technical assistance and Mara Ojeda for technical assistance with flow cytometry. We also wish to express our gratitude to the Wiener Lab for their contribution of ELISA reagents. Work on downregulation of *T. cruzi* bromodomains was funded by GSK (contract RCA3000030934 to RD and ES).

## Author Contributions

Conceptualization: V.B. E.S., P.S.R., A.R.P. and R.D.; Validation: E.S., V.P., P.S.R. and R.D.; Formal Analysis: V.B., V.G.P., J.A.C., F.P. and A.P.; Investigation: V.B., F.P., J.A.C., A.P. and V.P.; Resources: F.P., J.A.C.; Writing – Original Draft Preparation: V.B., V.G.P and E.S.; .Writing – Review & Editing: E.S., V.B., V.P., F.P., J.A.C., A.P., P.S.R., A.R.P. and R.D; Visualization; E.R.A. and G.F.; Supervision: E.S., P.S.R., R.D.; Funding Acquisition E.S. and R.D.

## References

1 PAHO / WHO - Chagas disease. 16 July 2025. Available: https://www.paho.org/en/topics/chagas-disease#3

2. Coura JR, Viñas PA. Chagas disease: a new worldwide challenge. Nature. 2010;465: S6–S7. doi:10.1038/nature09221

3. Chatelain E. Chagas disease research and development: Is there light at the end of the tunnel? Comput Struct Biotechnol J. 2017;15: 98–103. doi:10.1016/j.csbj.2016.12.002

4. Zuma AA, Dos Santos Barrias E, De Souza W. Basic Biology of Trypanosoma cruzi. Curr Pharm Des. 2021;27: 1671–1732. doi:10.2174/1381612826999201203213527

5. Cruz-Saavedra L, Vallejo GA, Guhl F, Ramírez JD. Transcriptomic changes across the life cycle of *Trypanosoma cruzi II*. PeerJ. 2020;8: e8947. doi:10.7717/peerj.8947

6. Teixeira SMR, DaRocha WD. Control of gene expression and genetic manipulation in the Trypanosomatidae. Genet Mol Res. 2003;2: 148–158.

7. Clayton C. Regulation of gene expression in trypanosomatids: living with polycistronic transcription. Open Biol. 2019;9: 190072. doi:10.1098/rsob.190072

8. Herreros-Cabello A, Callejas-Hernández F, Gironès N, Fresno M. Trypanosoma cruzi genome: Organization, multi-gene families, transcription, and biological implications. Genes. 2020;11: 1–26. doi:10.3390/genes11101196

9. Saha S. Histone Modifications and Other Facets of Epigenetic Regulation in Trypanosomatids: Leaving Their Mark. Li Z, Garsin DA, editors. mBio. 2020;11: e01079–20. doi:10.1128/mBio.01079-20

10. Fleck K, Nitz M, Jeffers V. “Reading” a new chapter in protozoan parasite transcriptional regulation. Kafsack BFC, editor. PLOS Pathog. 2021;17: e1010056. doi:10.1371/journal.ppat.1010056

11. Martínez-Calvillo S, Vizuet-De-Rueda JC, Florencio-Martínez LE, Manning-Cela RG, Figueroa-Angulo EE. Gene expression in trypanosomatid parasites. J Biomed Biotechnol. 2010;2010. doi:10.1155/2010/525241

12. Jacquet I, Paoli-Lombardo R, Vanelle P, Primas N. The epigenetic landscape of kinetoplastid parasites: From histone post-translational modifications to emerging therapeutic strategies. Bioorg Med Chem. 2025;131: 118377. doi:10.1016/j.bmc.2025.118377

13. Smith SG, Zhou M-M. The Bromodomain as the Acetyl-Lysine Binding Domain in Gene Transcription. In: Zhou M-M, editor. Histone Recognition. Cham: Springer International Publishing; 2015. pp. 1–26. doi:10.1007/978-3-319-18102-8_1

14. Shi J, Vakoc CR. The Mechanisms behind the Therapeutic Activity of BET Bromodomain Inhibition. Mol Cell. 2014;54: 728–736. doi:10.1016/j.molcel.2014.05.016

15. Pezza A, Tavernelli LE, Alonso VL, Perdomo V, Gabarro R, Prinjha R, et al. Essential Bromodomain *Tc* BDF2 as a Drug Target against Chagas Disease. ACS Infect Dis. 2022;8: 1062–1074. doi:10.1021/acsinfecdis.2c00057

16. Alonso VL, Ritagliati C, Cribb P, Serra EC. Construction of three new gateway® expression plasmids for Trypanosoma cruzi. Mem Inst Oswaldo Cruz. 2014;109: 1081–1085. doi:10.1590/0074-0276140238

17. Ritagliati C, Villanova GV, Alonso VL, Zuma AA, Cribb P, Motta MCM, et al. Glycosomal bromodomain factor 1 from Trypanosoma cruzi enhances trypomastigote cell infection and intracellular amastigote growth. Biochem J. 2016;473: 73–85. doi:10.1042/BJ20150986

18 Rodriguez Araya E, Merli ML, Cribb P, De Souza VC, Serra E. Deciphering Divergent Trypanosomatid Nuclear Complexes by Analyzing Interactomic Datasets with AlphaFold2 and Genetic Approaches. ACS Infect Dis. 2023;9: 1267–1282. doi:10.1021/acsinfecdis.3c00148

19. Allard S. NuA4, an essential transcription adaptor/histone H4 acetyltransferase complex containing Esa1p and the ATM-related cofactor Tra1p. EMBO J. 1999;18: 5108–5119. doi:10.1093/emboj/18.18.5108

20. Doyon Y, Selleck W, Lane WS, Tan S, Côté J. Structural and Functional Conservation of the NuA4 Histone Acetyltransferase Complex from Yeast to Humans. Mol Cell Biol. 2004;24: 1884–1896. doi:10.1128/MCB.24.5.1884-1896.2004

21. Squatrito M, Gorrini C, Amati B. Tip60 in DNA damage response and growth control: many tricks in one HAT. Trends Cell Biol. 2006;16: 433–442. doi:10.1016/j.tcb.2006.07.007

22. Hodges AJ, Plummer DA, Wyrick JJ. NuA4 acetyltransferase is required for efficient nucleotide excision repair in yeast. DNA Repair. 2019;73: 91–98. doi:10.1016/j.dnarep.2018.11.006

23. Staneva DP, Carloni R, Auchynnikava T, Tong P, Rappsilber J, Jeyaprakash AA, et al. A systematic analysis of *Trypanosoma brucei* chromatin factors identifies novel protein interaction networks associated with sites of transcription initiation and termination. Genome Res. 2021;31: 2138–2154. doi:10.1101/gr.275368.121

24. Devoucoux M, Roques C, Lachance C, Lashgari A, Joly-Beauparlant C, Jacquet K, et al. MRG Proteins Are Shared by Multiple Protein Complexes With Distinct Functions. Mol Cell Proteomics. 2022;21: 100253. doi:10.1016/j.mcpro.2022.100253

25. Rossetto D, Cramet M, Wang AY, Steunou A, Lacoste N, Schulze JM, et al. Eaf5/7/3 form a functionally independent NuA4 submodule linked to RNA polymerase II -coupled nucleosome recycling. EMBO J. 2014;33: 1397–1415. doi:10.15252/embj.201386433

26. Yamaguchi K, Nakagawa S, Furukawa Y. Understanding the role of BRD8 in human carcinogenesis. Cancer Sci. 2024;115: 2862–2870. doi:10.1111/cas.16263

27. Duraisingh MT, Horn D. Epigenetic Regulation of Virulence Gene Expression in Parasitic Protozoa. Cell Host Microbe. 2016;19: 629–640. doi:10.1016/j.chom.2016.04.020

28. Pezza A, Tavernelli LE, Alonso VL, Perdomo V, Gabarro R, Prinjha R, et al. Essential bromodomain TcBDF2 as a drug target against Chagas disease.

29. Kelly JM, Ward HM, Miles MA, Kendall G. A shuttle vector which facilitates the expression of transfected genes in Trypanosoma cruzi and Leishmania. Nucleic Acids Res. 1992 pp. 3963–3969. Available: https://academic.oup.com/nar/article/20/15/3963/2376634

30. Resende BC, Oliveira ACS, Guañabens ACP, Repolês BM, Santana V, Hiraiwa PM, et al. The Influence of Recombinational Processes to Induce Dormancy in Trypanosoma cruzi. Front Cell Infect Microbiol. 2020;10: 5. doi:10.3389/fcimb.2020.00005

31. Perdomo V, Boselli V, Manarin R, Serra E. Optimizing In Vitro Metacyclogenesis: Strain-Specific Variability in Trypanosoma cruzi Responses to Nutritional and pH Stress. 2025.

32. Marchini FK, De Godoy LMF, Rampazzo RCP, Pavoni DP, Probst CM, Gnad F, et al. Profiling the Trypanosoma cruzi Phosphoproteome. Li Z, editor. PLoS ONE. 2011;6: e25381. doi:10.1371/journal.pone.0025381

33. Chiurillo MA, Lander N, Bertolini MS, Storey M, Vercesi AE, Docampo R. Different Roles of Mitochondrial Calcium Uniporter Complex Subunits in Growth and Infectivity of *Trypanosoma cruzi*. Sibley LD, editor. mBio. 2017;8: e00574–17. doi:10.1128/mBio.00574-17

34. Teixeira TL, Chiurillo MA, Lander N, Rodrigues CC, Onofre TS, Ferreira ÉR, et al. Ablation of the P21 Gene of Trypanosoma cruzi Provides Evidence of P21 as a Mediator in the Control of Epimastigote and Intracellular Amastigote Replication. Front Cell Infect Microbiol. 2022;12: 799668. doi:10.3389/fcimb.2022.799668

35. Burle-Caldas GDA, Dos Santos NSA, De Castro JT, Mugge FLB, Grazielle-Silva V, Oliveira AER, et al. Disruption of Active Trans-Sialidase Genes Impairs Egress from Mammalian Host Cells and Generates Highly Attenuated Trypanosoma cruzi Parasites. Burleigh B, editor. mBio. 2022;13: e03478–21. doi:10.1128/mbio.03478-21

36. Alonso VL, Ritagliati C, Cribb P, Cricco JA, Serra EC. Overexpression of bromodomain factor 3 in Trypanosoma cruzi (TcBDF3) affects differentiation of the parasite and protects it against bromodomain inhibitors. FEBS J. 2016;283: 2051–2066. doi:10.1111/febs.13719

37. Sharma R, Peacock L, Gluenz E, Gull K, Gibson W, Carrington M. Asymmetric Cell Division as a Route to Reduction in Cell Length and Change in Cell Morphology in Trypanosomes. Protist. 2008;159: 137–151. doi:10.1016/j.protis.2007.07.004

38. Zuma AA, Dos Santos Barrias E, De Souza W. Basic Biology of Trypanosoma cruzi. Curr Pharm Des. 2021;27: 1671–1732. doi:10.2174/1381612826999201203213527

39. Matthews KR. The developmental cell biology of *Trypanosoma brucei*. J Cell Sci. 2005;118: 283–290. doi:10.1242/jcs.01649

40. Campbell PC, De Graffenried CL. Morphogenesis in Trypanosoma cruzi epimastigotes proceeds via a highly asymmetric cell division. Docampo R, editor. PLoS Negl Trop Dis. 2023;17: e0011731. doi:10.1371/journal.pntd.0011731

41. Vaughan S. Assembly of the flagellum and its role in cell morphogenesis in Trypanosoma brucei. Curr Opin Microbiol. 2010;13: 453–458. doi:10.1016/j.mib.2010.05.006

42. Campbell PC, De Graffenried CL. Morphogenesis in Trypanosoma cruzi epimastigotes proceeds via a highly asymmetric cell division. Docampo R, editor. PLoS Negl Trop Dis. 2023;17: e0011731. doi:10.1371/journal.pntd.0011731

43. Campbell PC, De Graffenried CL. Morphogenesis in Trypanosoma cruzi epimastigotes proceeds via a highly asymmetric cell division. Docampo R, editor. PLoS Negl Trop Dis. 2023;17: e0011731. doi:10.1371/journal.pntd.0011731

44. Hammarton TC, Engstler M, Mottram JC. The Trypanosoma brucei Cyclin, CYC2, Is Required for Cell Cycle Progression through G1 Phase and for Maintenance of Procyclic Form Cell Morphology. J Biol Chem. 2004;279: 24757–24764. doi:10.1074/jbc.M401276200

45. Zhao X, He Y, Zhang F, Aphasizheva I, Aphasizhev R, Zhang L. Comparative mitochondrial genome and transcriptome analyses reveal strain-specific features of RNA editing in *Trypanosoma brucei*. Nucleic Acids Res. 2025;53: gkaf661. doi:10.1093/nar/gkaf661

46. Sánchez-Valdéz FJ, Padilla A, Wang W, Orr D, Tarleton RL. Spontaneous dormancy protects Trypanosoma cruzi during extended drug exposure. eLife. 2018;7: e34039. doi:10.7554/eLife.34039

47. Barrett MP, Kyle DE, Sibley LD, Radke JB, Tarleton RL. Protozoan persister-like cells and drug treatment failure. Nat Rev Microbiol. 2019;17: 607–620. doi:10.1038/s41579-019-0238-x

48. Costa-Silva HM, Resende BC, Umaki ACS, Prado W, Da Silva MS, Virgílio S, et al. DNA Topoisomerase 3α Is Involved in Homologous Recombination Repair and Replication Stress Response in Trypanosoma cruzi. Front Cell Dev Biol. 2021;9: 633195w. doi:10.3389/fcell.2021.633195

49. Bunkofske ME, Sanchez-Valdez FJ, Tarleton RL. The importance of persistence and dormancy in Trypanosoma cruzi infection and Chagas disease. Curr Opin Microbiol. 2025;86: 102615. doi:10.1016/j.mib.2025.102615

50. Grünebast J, Lorenzen S, Zummack J, Clos J. Life Cycle Stage-Specific Accessibility of Leishmania donovani Chromatin at Transcription Start Regions. Gilbert JA, editor. mSystems. 2021;6: 10.1128/msystems.00628-21. doi:10.1128/msystems.00628-21

51. Siegel TN, Kawahara T, DeGrasse JA, Janzen CJ, Horn D, Cross GAM. Acetylation of histone H4K4 is cell cycle regulated and mediated by HAT3 in *Trypanosoma brucei*. Mol Microbiol. 2008;67: 762–771. doi:10.1111/j.1365-2958.2007.06079.x

52. Menezes APJ, Silber AM, Elias MC, Da Cunha JPC. Trypanosoma cruzi cell cycle progression exhibits minimal variation in histone PTMs with unique histone H4 acetylation pattern. J Proteomics. 2025;315: 105413. doi:10.1016/j.jprot.2025.105413

53. Santos CMBD, Ludwig A, Kessler RL, Rampazzo RDCP, Inoue AH, Krieger MA, et al. Trypanosoma cruzi transcriptome during axenic epimastigote growth curve. Mem Inst Oswaldo Cruz. 2018;113. doi:10.1590/0074-02760170404

54. Chávez S, Urbaniak MD, Benz C, Smircich P, Garat B, Sotelo-Silveira JR, et al. Extensive Translational Regulation through the Proliferative Transition of Trypanosoma cruzi Revealed by Multi-Omics. Phillips M, editor. mSphere. 2021;6: e00366–21. doi:10.1128/mSphere.00366-21

55. Chávez S, Eastman G, Smircich P, Becco LL, Oliveira-Rizzo C, Fort R, et al. Transcriptome-wide analysis of the Trypanosoma cruzi proliferative cycle identifies the periodically expressed mRNAs and their multiple levels of control. Te Pas MFW, editor. PLOS ONE. 2017;12: e0188441. doi:10.1371/journal.pone.0188441

56. Ramos TCP, Nunes VS, Nardelli SC, Dos Santos Pascoalino B, Moretti NS, Rocha AA, et al. Expression of non-acetylatable lysines 10 and 14 of histone H4 impairs transcription and replication in Trypanosoma cruzi. Mol Biochem Parasitol. 2015;204: 1–10. doi:10.1016/j.molbiopara.2015.11.001

57. Filippakopoulos P, Picaud S, Mangos M, Keates T, Lambert J-P, Barsyte-Lovejoy D, et al. Histone Recognition and Large-Scale Structural Analysis of the Human Bromodomain Family. Cell. 2012;149: 214–231. doi:10.1016/j.cell.2012.02.013

58. Lander N, Li ZH, Niyogi S, Docampo R. CRISPR/Cas9-induced disruption of paraflagellar rod protein 1 and 2 genes in Trypanosoma cruzi reveals their role in flagellar attachment. mBio. 2015;6: 1–12. doi:10.1128/mBio.01012-15

59. Lander N, Chiurillo MA, Docampo R. Genome editing by CRISPR/Cas9 in trypanosoma cruzi. Methods Mol Biol. 2019;1955: 61–76. doi:10.1007/978-1-4939-9148-8_5

60. Alonso VL. Ultrastructure Expansion Microscopy (U-ExM) in Trypanosoma cruzi: localization of tubulin isoforms and isotypes. Parasitol Res. 2022;121: 3019–3024. doi:10.1007/s00436-022-07619-z

61. De Hernández MA, Martinez Peralta G, Vena R, Alonso VL. Ultrastructural Expansion Microscopy in Three In Vitro Life Cycle Stages of Trypanosoma cruzi. J Vis Exp. 2023; 65381. doi:10.3791/65381

62. Rodrigues JPF, Souza Onofre T, Barbosa BC, Ferreira ÉR, Bonfim-Melo A, Yoshida N. Host cell protein LAMP-2 is the receptor for *Trypanosoma cruzi* surface molecule gp82 that mediates invasion. Cell Microbiol. 2019;21: e13003. doi:10.1111/cmi.13003

63. Andrade LO, Andrews NW. The Trypanosoma cruzi–host-cell interplay: location, invasion, retention. Nat Rev Microbiol. 2005;3: 819–823. doi:10.1038/nrmicro1249

64. Romano PS, Cueto JA, Casassa AF, Vanrell MC, Gottlieb RA, Colombo MI. Molecular and cellular mechanisms involved in the *Trypanosoma cruzi* /host cell interplay. IUBMB Life. 2012;64: 387–396. doi:10.1002/iub.1019

65. Brener Z. Therapeutic activity and criterion of cure on mice experimentally infected with Trypanosoma cruzi. Rev Inst Med Trop Sao Paulo. 1962;4: 389–396.

66. Cummings KL, Tarleton RL. Rapid quantitation of Trypanosoma cruzi in host tissue by real-time PCR. Mol Biochem Parasitol. 2003;129: 53–59. doi:10.1016/S0166-6851(03)00093-8

67. Duffy T, Bisio M, Altcheh J, Burgos JM, Diez M, Levin MJ, et al. Accurate Real-Time PCR Strategy for Monitoring Bloodstream Parasitic Loads in Chagas Disease Patients. Rodriguez A, editor. PLoS Negl Trop Dis. 2009;3: e419. doi:10.1371/journal.pntd.0000419

