## Supplementary material for "BDF6 deficiency severely compromises intracellular amastigote development and infectivity of *Trypanosoma cruzi*": Table S1, Fig. S1, Fig S2, Fig S3, FingS4, Fig S5

1. Instituto de Biología Molecular y Celular de Rosario, IBR-CONICET, Rosario, Santa Fe, Argentina. 2. Facultad de Ciencias Bioquímicas y Farmacéuticas, Universidad Nacional de Rosario, Rosario, Santa Fe, Argentina. 3. CONICET, Rosario, Argentina. 4. Instituto de Inmunología Clínica y Experimental de Rosario, IDICER-CONICET, Rosario, Santa Fe, Argentina. 5. Laboratorio de Biología de *Trypanosoma cruzi* y la célula hospedadora, Instituto de Histología y Embriología (IHEM) "Dr. Mario H. Burgos", Mendoza, Mendoza, Argentina. 6. Facultad de Ciencias Médicas, Universidad Nacional de Cuyo, Mendoza, Mendoza, Argentina. 7. Center for Tropical and Emerging Global Diseases and Department of Cellular Biology, University of Georgia, Athens, GA, USA. 8. Centro de Investigación y Producción de Reactivos Biológicos (CIPReB), Facultad de Ciencias Médicas, Universidad Nacional de Rosario, Rosario, Santa Fe, Argentina.

### Supplemental Material

Table S1. Description of the oligonucleotides. Oligonucleotides and ultramers used for the generation of the *TcBDF6* $^{-/-}$ strain

| Oligonucleotides | Sequence |
|---|---|
| **sgRNAFw** | 5′-ATCGGATCC**GCGGCGGGAAGATTTACTGC**GTTTTAGAGCTAGAAATAGC-3′ |
| **sgRNARv** | 5′-CAGTGGATCCAAAAAAGCACCGACTCGGTG-3′ |
| **Val_BDF6_KO_Fw** | 5´- GCTGCTGCTTGCTTGTTTGTGAG-3´ |
| **Val_BDF6_KO_Rv** | 5´-TCTTCTTCGATTTTGATTTTTTGATC-3´ |
| **TcBDF6_MutCas9** | 5´-AAGGATCCATGTATCCGTATGATGTGCCGGATTATGCTCGACGAGAGGACTTGCTACAGGTGTTGCTGGTC-3´ |
| **TcBDF6_EcoRV** | 5´-AAAGATATCTCATGCACCACGCAAATGC -3´ |
| **TCZ-F** | 5´ -GCTCTTGCCCA CAAGGGTGC-3´ |
| **TCZ-R** | 5´-CCAAGCAGCGGATAGTTCAGG-3´ |
| **Ultramers** | **Sequence** |
| ***Tc*BDF6_BSD_KO Fw** | 5′-AACACATACACACACAGAGAGAGAAAGAGAGAGAAAGGGAGAGAAAGAGAAAAGGGGAGTTGTGTCGGCCGGAGCGTGTAGAAAAAGATGCGGCGGGAAGATGGCCAAGCCTTTGTCTCA-3′ |
| ***Tc*BDF6_BSD_KO Rv** | 5′-ACCTTCCCCCACACCTTCGACGGGACGGTCCAAACATCAAGTCACAAAAAGGGGCGTTGCACATGCATCCATCATCCGACGCCTTAGCCCTCCCACACATAAC-3′ |

**Fig S1. Genotyping of *T. cruzi* mutant strains by PCR.** PCR amplification was performed using oligonucleotides flanking the insertion site of the blasticidin resistance cassette (Val_KO_BDF6_Dm28c oligonucleotides). The WT strain shows a single band of 1391 bp, corresponding to the endogenous *tcbdf6* allele. The heterozygous *TcBDF6* $^{-/+}$ strain shows two bands of 1391 bp and 1700 bp, corresponding to the disrupted allele with the resistance cassette. The homozygous *TcBDF6* $^{-/-}$ clones (Clone A and Clone B) display the single band of 1700 bp. The add back strain *TcBDF6* $^{-/-}$ pBDF6 shows the same pattern as the homozygous KOs, confirming that complementation was performed in the *TcBDF6* $^{-/-}$ background.

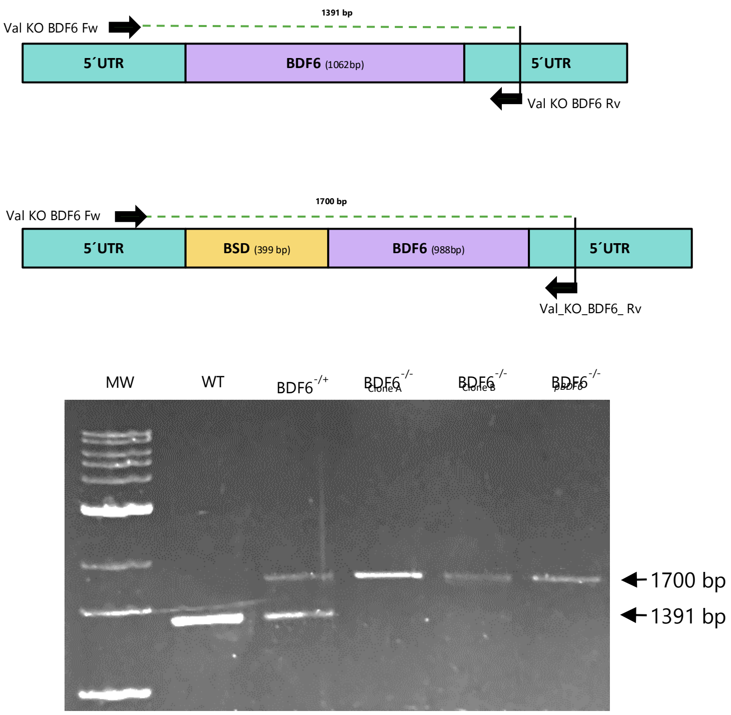


**Fig S2. Epimastigote growth, infectivity, and intracellular differentiation with bromodomain mutant add back BDF6$^{-/-}_{S97A}$.** (A) Growth curves of epimastigotes from WT, *TcBDF6* $^{-/-}$, add back (*TcBDF6* $^{-/-}$pBDF6), and bromodomain mutant add back (*TcBDF6* $^{-/-}$pBDF6$_{S97A}$) parasites cultured over six days. Data represent mean ± SD. (B) Representative fluorescence microscopy images of infected host cells showing intracellular amastigotes for the indicated parasite strains. Nuclei and kinetoplasts are visualized by DAPI staining. (C) Percentage of infected cells quantified for each parasite strain. (D) Number of amastigotes per infected cell. Data in (C) and (D) represent mean ± SD from at least three independent experiments. Statistical significance was determined using Tukey's test, with *$p < 0.05$, **$p < 0.01$, ***$p < 0.001$

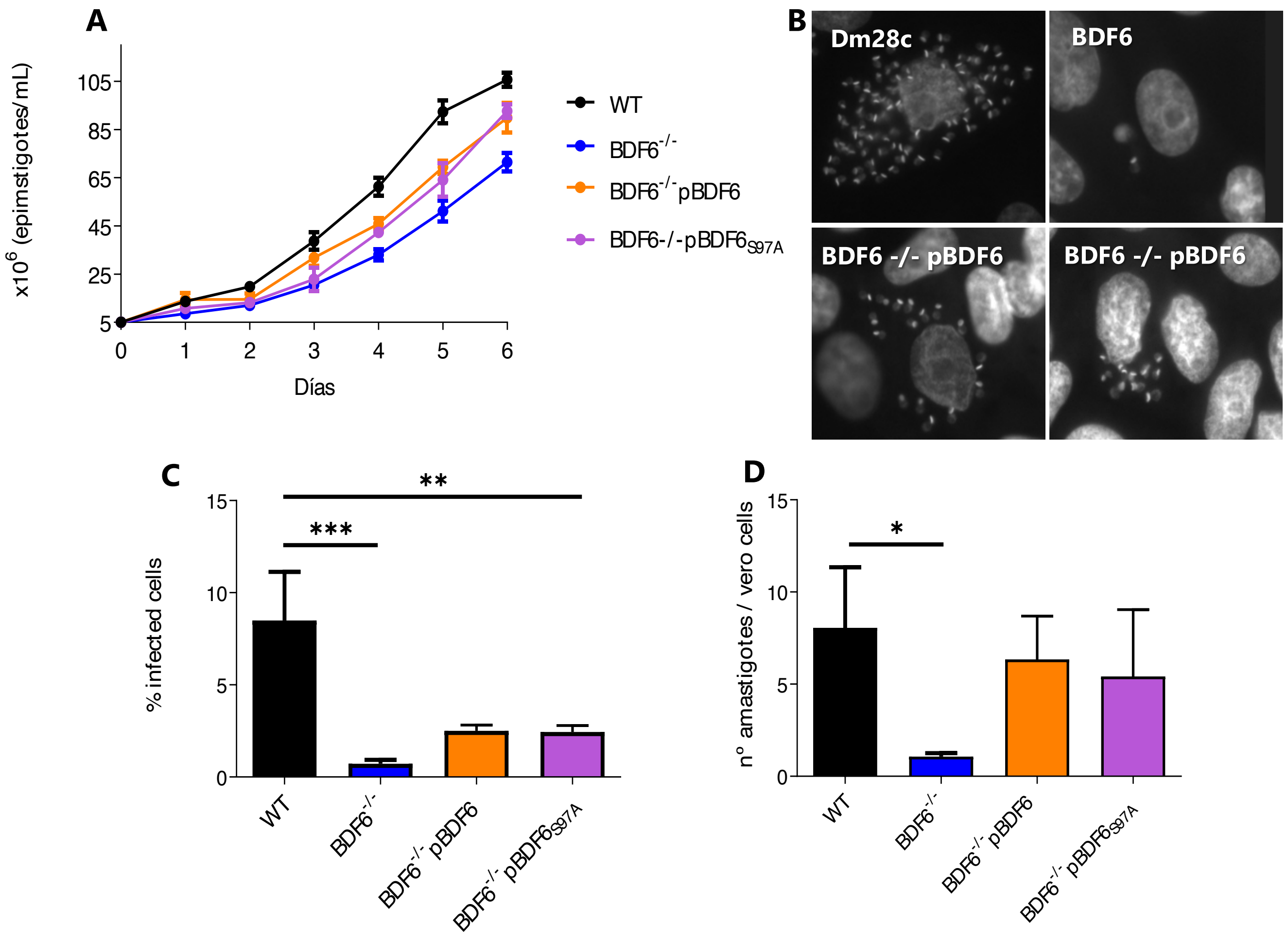


**Fig S3. Quantification of epimastigotes cell surface area.** Violin plot representing the area (in μm$^2$) of epimastigotes forms from WT, heterozygous (*TcBDF6* $^{-/+}$), KO (*TcBDF6* $^{-/-}$), and add back (*TcBDF6* $^{-/-}$ pBDF6) strains. Cell surface areas were quantified from microscope images using ImageJ. Violins indicate median and interquartile ranges.

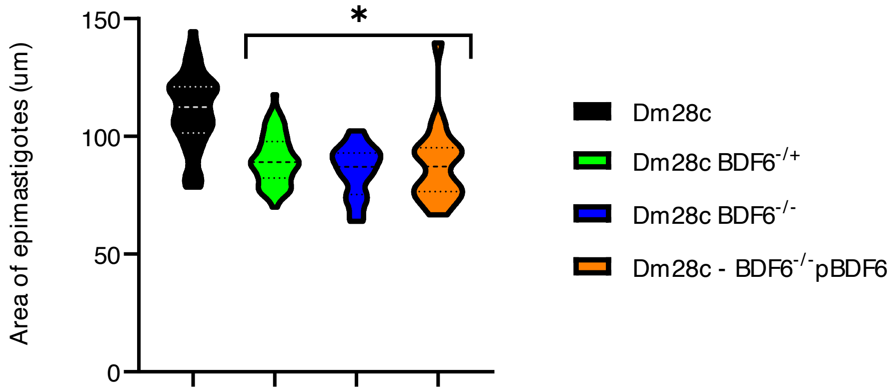

**Fig S4. Deletion of *TcBDF6* enhances *T. cruzi* metacyclogenesis.** Percentage of metacyclic trypomastigotes formed during *in vitro* metacyclogenesis was quantified in WT, *TcBDF6* $^{-/+}$ heterozygous, *TcBDF6* $^{-/-}$ KO clones (A and B), and add back strain *TcBDF6* $^{-/-}$pBDF6. Data represent mean ± SD of three independent experiments.

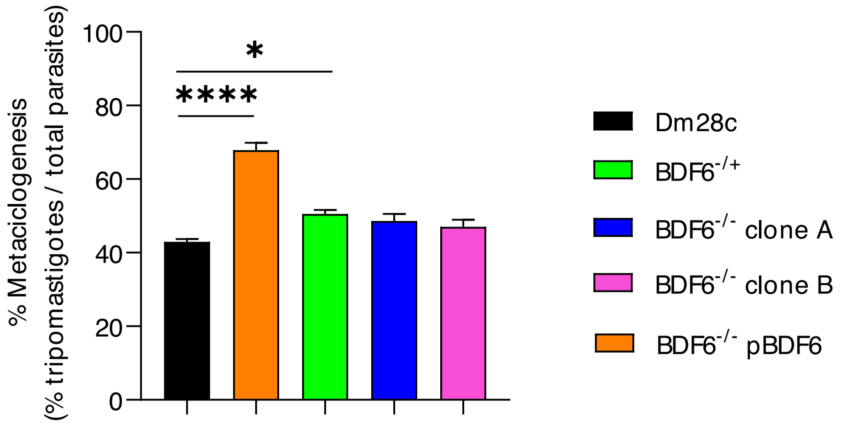


**Fig S5. Anti-*T. cruzi* IgG response**. Serum IgG levels were measured by ELISA at 53 days p.i. in C57BL/6 mice infected with metacyclic trypomastigotes WT, *TcBDF6* $^{-/+}$, *TcBDF6* $^{-/-}$, or the add back strain *TcBDF6* $^{-/-}$pBDF6 and treated from day 31 onwards with 3 doses of cyclophosphamide. A group of uninfected animals subject to the same protocol was included as a negative control. Bars represent mean ± SD. Results represent two independent experiments with 3–5 animals per group. Statistical comparisons between groups were performed using non-parametric tests. Significance levels are indicated as follows: *p < 0.05, **p < 0.01, ***p < 0.001 (between specified groups); #p < 0.05, ##p < 0.01 versus all other groups.

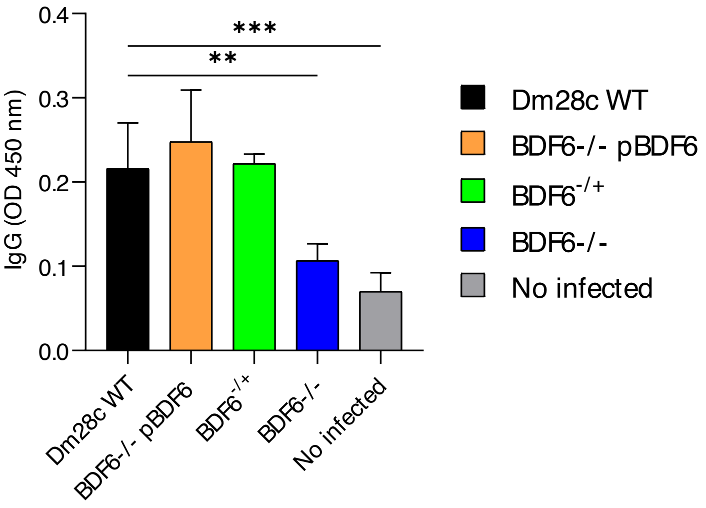
